# Aerobic growth physiology of *Saccharomyces cerevisiae* on sucrose is strain-dependent

**DOI:** 10.1101/2021.02.25.432870

**Authors:** Carla Inês Soares Rodrigues, Aljoscha Wahl, Andreas K. Gombert

## Abstract

Present knowledge on the quantitative aerobic physiology of the yeast *Saccharomyces cerevisiae* during growth on sucrose as sole carbon and energy source is limited to either adapted cells or to the model laboratory strain CEN.PK113-7D. To broaden our understanding of this matter and open novel opportunities for sucrose-based biotechnological processes, we characterized three strains, with distinct backgrounds, during aerobic batch bioreactor cultivations. Our results reveal that sucrose metabolism in *S. cerevisiae* is a strain-specific trait. Each strain displayed a distinct extracellular hexose concentration and invertase activity profiles. Especially, the inferior maximum specific growth rate (0.21 h^−1^) of the CEN.PK113-7D strain, with respect to that of strains UFMG-CM-Y259 (0.37 h^−1^) and JP1 (0.32 h^−1^), could be associated to its low invertase activity (0.04 to 0.09 U mg_DM_^−1^). Moreover, comparative experiments with glucose or fructose alone, or in combination, suggest mixed mechanisms of sucrose utilization by the industrial strain JP1, and points out the remarkable ability of the wild isolate UFMG-CM-259 to grow faster on sucrose than on glucose in a well-controlled cultivation system. This work hints to a series of metabolic traits that can be exploited to increase sucrose catabolic rates and bioprocess efficiency.

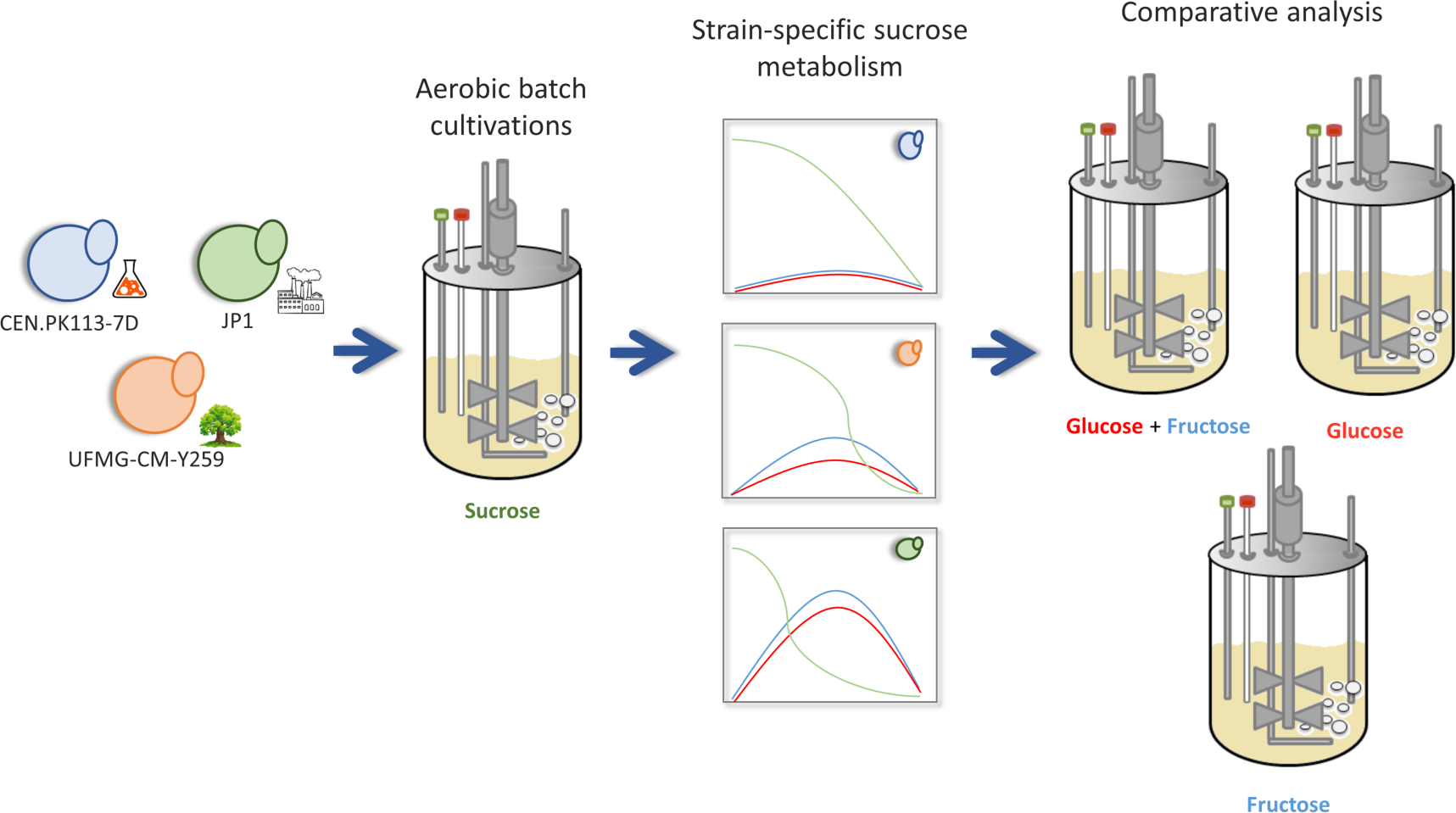

## INTRODUCTION

Sucrose has long been used in the food industry as a substrate for the production of bakery goods and beverages. In the last decades, however, it has also been considered a valuable feedstock for the replacement of petrochemical derived materials, due to its low market price (Polat and Linhardt 2001) (Marques et al. 2016). Especially, sucrose feedstocks have a high land efficiency and vast availability of sugar-rich crops. Furthermore, sucrose does not require any pretreatment prior to its use in industrial fermentation. Whilst fuel ethanol is by far the prime product of the nonfood sucrose-based industry (OECD-FAO 2018), with an annual production of 33.1 billion liters in Brazil alone in 2019 (Brazilian Sugarcane Industry Union (Unica)), the potential use of this disaccharide goes beyond fuel manufacturing, with value-added chemicals such as citric acid (Förster et al. 2007; Show et al. 2015), lactic acid (Lunelli et al. 2010; Wang et al. 2012) and farnesene (E4tech et al. 2015) also being successfully commercialized. Several additional chemical intermediates can be produced using sucrose as substrate, including 5-hydroxymethylfurfural (Peters et al. 2010; E4tech et al. 2015), 1-2-propylene glycol (Peters et al. 2010; Rosales-Calderon and Arantes 2019), acrylic acid (E4tech et al. 2015), and succinic acid (Chan et al. 2012; Jiang et al. 2014; E4tech et al. 2015; Rosales-Calderon and Arantes 2019).

The yeast *Saccharomyces cerevisiae*, the primary workhorse of the biotechnology industry (Steensels et al. 2014; Steensels and Verstrepen 2014; Kavšček et al. 2015), naturally metabolizes sucrose either through its hydrolysis in the periplasmic space or through direct uptake via active transport of the disaccharide and its hydrolysis in the cytosol (Badotti et al. 2008) (**Figure 1**). In the first mechanism, *S. cerevisiae* needs to express an invertase-encoding gene (*SUC2*, the most common one) and secrete the protein to the periplasmic space after oligomerization and post-translational modification (Carlson and Botstein 1982). The hydrolysis’ products, namely glucose and fructose, can enter the cells by facilitated diffusion. In the other mechanism, sucrose is directly transported to the cytosol via a proton-symport mechanism mediated by the high-affinity (K_M_ = 7.9 ± 0.8 mM) transporter Agt1p or the low-affinity (K_M_ = 120 ± 20 mM) transporters encoded by *MALx1* genes (x denotes the locus number) (Stambuk et al. 2000; Basso et al. 2011). In this case, ATP is invested to expel the imported proton with a stoichiometry of 1:1, in order to keep cell’s homeostasis and the proton motive force across the plasma membrane. Consequently, cells are expected to achieve a higher glycolytic rate to compensate for the lower energy efficiency in the overall metabolic process.

**Figure 1.**
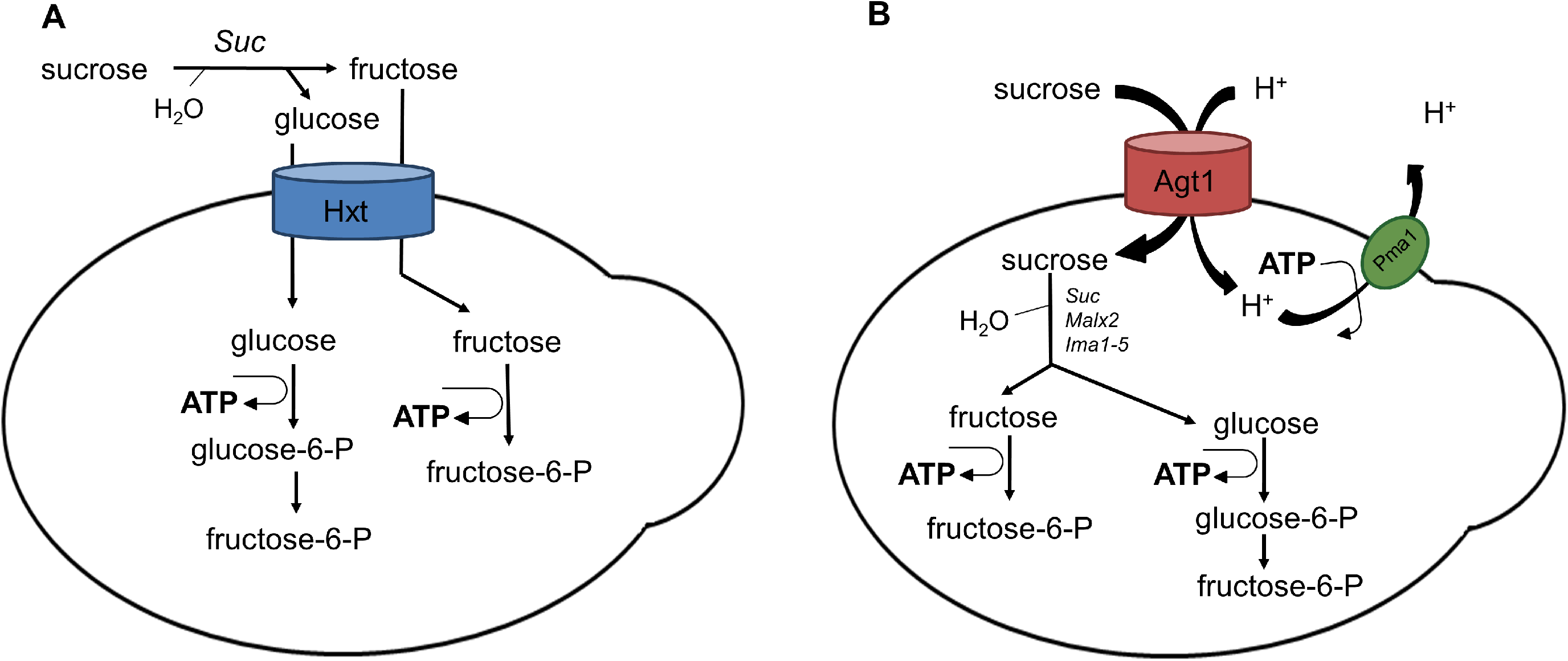
Schematic illustration of the two mechanisms of sucrose utilization in *S. cerevisiae*: A) periplasmic hydrolysis by invertase (Suc) and passive transport of glucose and fructose into the cells; B) active sucrose transport via the **A**lpha-**G**lucoside **T**ransporter (Agt1) and ATP-dependent export of H^+^ via the **P**lasma **M**embrane **A**TPase (Pma1).

Aerobic sucrose metabolism by *S. cerevisiae* has been shown to be fast, with maximum specific growth rates ranging from 0.38 to 0.57 h^−1^ (van Dijken et al. 2000; Beato et al. 2016), depending on the strain, conditions and analytical methods employed. In spite of this successful relationship between sucrose and *S. cerevisiae* (Marques et al. 2016), little attention has been given to the specific effects of this carbon source on the quantitative aerobic physiology of this yeast. The few studies involving a comprehensive quantitative aerobic physiological analysis were performed with strains pre-evolved on sucrose for 200 to 250 generations (Orlowski and Barford 1991; Barford et al. 1995; Mwesigye and Barford 1996), or with the model laboratory strain CEN.PK113-7D (Herwig et al. 2001), which limits our understanding of relevant phenomena. On the one hand, the use of pre-adapted strains changes their initial/natural physiology. On the other hand, the results published on a non-adapted strain refer to one particular laboratory strain, meaning that we still do not have a good overview of the physiology of this yeast species during growth on sucrose and how this eventually varies among different strains. Relevant biological questions remain to be elucidated: does the strain background influence *S. cerevisiae*’s physiology on sucrose?; Would a non-adapted fast sucrose-growing strain rely on direct uptake as a preferred natural mechanism of sucrose utilization?; How does growth on sucrose compare to growth on glucose? A better understanding of *S. cerevisiae*’s growth on sucrose will not only aid in answering those questions but also open novel opportunities for the development of strain improvement strategies to enhance sucrose-based industrial bioprocesses, in terms of the yields, productivities, and/or titers required for the production of market goods in a competitive manner.

Here, we present the quantitative physiology of *S. cerevisiae* grown on sucrose as the sole carbon and energy source in aerobic batch bioreactor cultivations. To access the effects of the strain background on sucrose physiology, the experiments were performed with one laboratory (CEN.PK113-7D), one industrial (JP1) and one wild isolate (UFMG-CM-Y259) strain. While the CEN.PK113-7D strain serves as a reference for physiological studies (Van Dijken et al. 2000), JP1 and UFMG-CM-Y259 were chosen due to their different behaviors on sucrose (Beato et al. 2016). JP1 was isolated from fermentors used to produce fuel ethanol from sugarcane in the Northeast of Brazil and is a relatively thermotolerant strain (da Silva Filho et al. 2005; Della-bianca and Gombert 2013), whereas UFMG-CM-Y259 presented the highest maximum specific growth rate (µ_MAX_) on sucrose in a screening test made with ~20 different *S. cerevisiae* strains. Its µ_MAX_ on sucrose was also ~20% higher than the corresponding value on glucose (Beato et al. 2016).

Additionally, in order to investigate the mechanisms underlying the diverging physiologies observed during growth on sucrose, cultivations were also performed on an equimolar mixture of glucose and fructose and on each one of these two monosaccharides separately.

## MATERIAL AND METHODOLOGY

### Yeast strains, preservation and pre-cultures

The *Saccharomyces cerevisiae* strains studied in this work comprise an indigenous strain, named UFMG-CM-Y259, isolated from barks of the tree *Quercus rubra* (Northern Red Oak) located in Santuário do Caraça (Minas Gerais, Brazil) within the Atlantic Forest biome (Beato et al. 2016), kindly provided by Dr. Carlos A. Rosa (Federal University of Minas Gerais, Belo Horizonte, Brazil); JP1, a Brazilian fuel ethanol industrial strain isolated from the Japungu Agroindustrial sugarcane-based distillery located in Northeastern Brazil (da Silva Filho et al. 2005), kindly provided by Dr. Marcos Morais Jr. (Federal University of Pernambuco, Recife, Brazil); and the laboratory strain CEN.PK113-7D (kindly provided by Dr. Petter Kötter, EUROSCARF, Germany), which is largely employed in physiological studies by the scientific community (van Dijken et al. 2000).

Stock cultures were prepared by growing cells until stationary phase in 500-ml Erlenmeyer flasks containing 100 ml YPD medium (per liter: 10.0 g **Y**east extract, 20.0 g **P**eptone and 20.0 g **D**extrose/Glucose), in an incubator shaker (Certomat BS-1, Braun Biotech International, Berlin, Germany) operating at 30 °C and 200 rpm for 24 h. 20% (v/v, final concentration) glycerol was added and 1 ml aliquots were stored in 2-ml cryogenic vials in an ultra-freezer (CryoCube HEF, model F570h-86, Eppendorf, Hamburg, Germany) at −80 °C until further use.

The pre-culture medium was prepared according to (Verduyn et al. 1992) with a few modifications. The medium consisted of (per liter): 3.0 g KH_2_PO_4_, 6.6 g K_2_SO_4_, 0.5 g MgSO_4_.7H_2_O, 2.3 g urea, 1 ml trace elements solution, 1 ml vitamins solution and 10 g carbon source (sucrose, glucose, fructose or an equimolar mixture of glucose and fructose). The initial pH of the pre-culture medium was adjusted to 6.0 using 2 mol/l KOH. Sterilization of the pre-culture medium occurred by filtration through 0.22 µm pore membranes (Millex-GV, Merck Millipore, Massachusetts, USA).

To prepare the inoculum for bioreactor cultivations, the content of one stock cryogenic vial was centrifuged at 867 *g* for 4 min, and cells were transferred to a 500-ml Erlenmeyer flask containing 100 ml of the pre-culture medium. The pre-inoculum was left in a shaker (Certomat BS-1, Braun Biotech International) set at 30 °C and 200 rpm for 24 h preceding the direct transfer of 1 ml of its content to another shake-flask with fresh pre-culture medium. After a second round of 24 h of growth in a shaker, operating at the same settings as before, an aliquot sufficient to start the bioreactor batch cultivation with an optical density of 0.2 at 600 nm was collected and washed. For the washing procedure, cells were centrifuged at room temperature and 3500 *g* for 3 min, the supernatant was discarded and fresh pre-culture medium was added to the cell pellet. This was then vortexed and centrifuged once again. At last, the cells were resuspended with cultivation medium and transferred to a proper flask that allows for an aseptic transfer to the bioreactor.

### Bioreactor batch cultivations

A synthetic medium formulated as described in (Verduyn et al. 1992) was used in all bioreactor batch cultivations. The medium contains per liter: 3.0 g KH_2_PO_4_, 5.0 g (NH_4_)_2_SO_4_, 0.5 g MgSO_4_.7H_2_O, 1 ml trace elements solution, 1 ml vitamins solution and 20 g (or equivalent in g_GLCeq_) carbon and energy source — sucrose, glucose, fructose or an equimolar mixture of glucose and fructose. The initial pH was adjusted to 5.0 using of 4 mol/l NaOH. Sterilization of the medium occurred by filtration through 0.22 µm pore membranes (Millex-GV, Merck Millipore, Massachusetts, USA).

A 2-liter bioreactor (Applikon Biotechnology B.V., Delft, The Netherlands), with a working volume of 1.2 l was used throughout this work. Cells were grown at 30°C and 800 rpm stirring speed. Aeration occurred with compressed air at 0.5 l min^−1^ flow rate using a mass flow controller (Model 58505, Brooks Instrument, Hatfield, USA). The pH of the broth was controlled at 5.0 by automatic addition of a 0.5 mol/l KOH solution. A 10% (v/v) antifoam C emulsion (Sigma-Aldrich, Missouri, USA) was added manually to the broth upon necessity. Aliquots were collected manually at different sampling times to be analysed for extracellular metabolites concentrations, dry cell mass and invertase activity. For each collected sample, the exact mass withdrawn was determined.

The gas flowing out of the bioreactor had its CO_2_ and O_2_ molar fractions determined using a gas analyzer device (Rosemount NGA 2000, Emerson Electric Co., Ferguson, USA). Determination of O_2_ molar fraction was performed through a paramagnetic detector, while an infrared detector was used to determine CO_2_ molar fraction. Bioreactor volumetric rates were calculated taking into account variations in pressure and volume (e.g. due to sampling, base and antifoam addition).

Cultivations were finished when a decrease in the CO_2_ molar fraction in the off-gas was observed.

## Analytical Methods

### Dry cell mass concentration

Cells were harvested and filtered through a 0.45 µm nitrocellulose membrane (SO-Pak filters, HAWP047S0 – Merck Millipore, Massachusetts, USA) that had been previously dried and weighed (m_1_). The cell pellet was washed twice with demineralized water. The filter containing the pellet was dried in an oven at 70 °C for 48 h and then placed in a desiccator to cool down prior to being weighed (m_2_). The cell dry mass (X_DM_) was calculated by dividing the difference between the filter’s mass after and before filtration by the sample volume filtered (V); X_DM_ = (m_2_– m_1_)/V. The result is expressed in g_DM_.l^−1^.

### Concentration of extracellular metabolites

Sampling for extracellular metabolites followed the procedure described elsewhere (Mashego et al. 2003). Briefly, a defined volume of broth was rapidly collected in a syringe containing a calculated amount of cold metal beads (−20°C) - enough to cool the collected broth to 1°C, and filtered through a 0.45 µm PVFD membrane (Millex - HV, Merck Millipore) directly into a tube. All sample tubes containing filtrate were immediately placed on ice and stored at - 80°C until analysis.

The concentrations of residual sugars in the filtrate - sucrose, glucose and fructose - were determined either enzymatically using the K-SURFG kit (Megazyme, Bray, Ireland), following the manufacturer’s instructions, or by means of ion chromatography coupled with pulsed electrochemical detection at gold electrode (CarboQuad pulse, AgCl reference). The chromatography system was a Dionex ICS – 5000 HPIC system with AS-AP sampler, SP pump (Thermal Scientific, Waltham, USA) equipped with a Carbopac PA-20 3×150 mm column and Aminotrap 3×30 mm precolumn at 30°C, eluted at 0.5 ml min^−1^ with 5% NaOH 200 mM for 15 min, followed by a cleaning step with 20% sodium acetate solution 0.5 M in 200 mM NaOH for 5 min, and reequilibration for 15 min with Milli-Q water (eluents C and D). The temperature of the detector was kept at 15°C.

The concentrations of ethanol, glycerol and organic acids (lactate, succinate and acetate) were always determined by high performance liquid chromatography (HPLC) using a Bio-Rad Aminex HPX-87H column (Bio-Rad, USA), kept at 60°C and eluted with highly diluted phosphoric acid (60 ml min^−1^) at a pH between 2 and 3, which was preheated before use. The samples were injected using an autosampler (Waters 717, USA). Detection of organic acids was performed via a UV detector (Waters 2489), while detection of ethanol or glycerol was performed using a refraction index detector (Waters 2414, USA). During cultivations without sucrose, residual glucose and fructose were determined in the same HPLC run used for the other metabolites, and the detection of these hexoses was performed using a refraction index detector (Waters 2414, USA). HPLC data were processed using Empower software (Waters Corporation, Milford, USA).

### Extracellular invertase activity

Extracellular (or periplasmic) invertase activity was determined according to the approach described in (Silveira et al. 1996), with a few modifications. Briefly, cells were centrifuged and resuspended in distilled water such that a 20 g_DM_.l^−1^ suspension was obtained. Next, the cells were treated with Succinate-Tris buffer (pH 5.0) containing sodium fluoride, which is an inhibitor of enolase. A sucrose solution was added to the reaction mixture and the glucose formed at 30°C due to disaccharide hydrolysis was measured using an enzymatic kit (R-Biopharm AG, Darmstadt, Germany). Invertase activity was reported as µmol of glucose produced per minute per milligram of cell dry mass (U mg_DM_^−1^).

### Calculation of physiological parameters

Prior to calculating physiological parameters, the experimental data points were treated as follows. Extracellular metabolites concentrations and CO_2_ and O_2_ amounts were plotted against time and a polynomial was fitted to the data. Concentrations or amounts were calculated for each time point taking the polynomial equation. Glucose and fructose concentrations during experiments performed with either sucrose or an equimolar mixture of glucose and fructose were not adjusted to avoid concealing their actual consumption trend. Only data points within the exponential growth phase (EGP) were considered for the calculation of physiological parameters.

The specific growth rate during the EGP (µ_MAX_) was calculated as the slope of the straight line adjusted to the linear region of an ln(X_DM_) *versus* time plot. The time span corresponding to this linear region was considered to be the EGP. Biomass (Y_X/S_) and product yield (Y_P/S_) on substrate, except for CO_2_, were calculated as the absolute value of the slope of a biomass concentration (X_DM_) *versus* substrate concentration (S) plot, and of a product concentration (P) *versus* substrate concentration (S) plot, respectively. CO_2_ yield on substrate was derived from the absolute value of the slope of an integrated CO_2_ amount versus substrate amount plot. For cultivations with sucrose or a mixture of glucose and fructose, substrate concentration (S) was determined as the sum of the concentrations of all carbohydrates in g_GLGeq_ l^−1^. For that, 1 g of sucrose was considered to be equivalent to 1.0526 g of hexose.

The maximum specific substrate consumption rate (during the EGP) was calculated as the ratio between the maximum specific growth rate and the biomass yield on substrate; q_S,MAX_ = µ_MAX_/Y_X/S_.

The maximum specific product formation rate (q_P_) was calculated taking the ratio between the desired yield on substrate and biomass yield on substrate, and multiplying the result by the maximum specific growth rate; q_P,MAX_ = µ_MAX_*Y_P/S_/Y_X/S_

For determining the specific oxygen consumption rate (q_O2_), the integrated amount in mmol of oxygen consumed per gram of substrate consumed at each sampling time was first obtained by the absolute slope of the integrated O_2_ *versus* substrate amount plot. Following, this value was divided by the biomass yield on substrate and multiplied by the maximum specific growth rate.

### Calculation of the percentage of fermented sugar during the EGP

The specific rate of ethanol formation (q_ETH_) was used to calculate the specific rate of CO_2_ formed due to fermentation (q_CO2_ferm_) in the EGP. This value was divided by 2 and considered to be the specific rate of sugar that was fermented (e.g. q_GLC_ferm_), as a result of the stoichiometry of ethanolic fermentation (C_6_H_12_O_6_ → 2 C_2_H_6_O + 2 CO_2_). This value, in turn, was divided by the (total) maximum specific substrate consumption rate (e.g. q_GLC,MAX_) and multiplied by 100, resulting in the % fermented sugar.

### Sequencing of the *S. cerevisiae* JP1 and UFMG-CM-Y259 genomic DNA

The *S. cerevisiae* JP1 and Y259 strains were sequenced at Serviço Nacional de Aprendizagem Industrial (SENAI), Rio de Janeiro, Brazil. DNA extraction was performed using the Wizard® Genomic DNA Purification kit (Promega) and sequencing libraries were prepared using the Nextera DNA Flex kit (Illumina), always following the manufacturer’s guidelines. Sequencing was carried out on an Illumina NextSeq 550 machine, using a High Output cartridge, producing 300 bp reads (2x 151 bp paired-end). Coverage (relative to a 12 Mbp haploid genome) was 427x for the UFMG-CM-Y259 strain and 174x for the JP1 strain. The reads can be accessed using the BioProject ID PRJNA699867 at NCBI.

### Bioinformatics pipeline

The fastq files containing the sequencing results were analyzed using a pipeline recommended by the Research Computing Faculty of Applied Sciences (RCFAS), Harvard University, which included: 1) trimming with NGmerge (Gaspar, 2018), 2) alignment against the *S. cerevisiae* S288c reference genome (yeastgenome.org, release R64) with Bwa (Li and Durbin, 2009), 3) conversion of .sam to .bam files and validation of the final .bam file with Picard <u>(ht</u>t<u>p://broadinstitute.github.io/picard),</u> 4) haplotype calling with GATK HaplotypeCaller (McKenna et al, 2010), 5) databasing with GATK GenomicsDBImport, and 6) genotyping with GATK GenotypeGVCFs. The .bam files generated with Picard were used to manually inspect the desired variants using the Integrated Genome Viewer (IGV, Robinson et al. 2011). The final .vcf files generated with GATK were used to annotate variants using SnpEff (Cingolani et al. 2012).

## RESULTS AND DISCUSSION

### *S. cerevisiae* strains from different environments display different physiologies on sucrose

The physiology of the *S. cerevisiae* strains CEN.PK113-7D, UFMG-CM-Y259 and JP1 during growth on sucrose was assessed in controlled batch aerobic bioreactor cultivations (**Figures 2 and 3; Table 1; Supplementary Figures S1-S5, Supplementary Tables S1-S3**). The laboratory strain CEN.PK113-7D displayed the lowest maximum specific growth rate on sucrose (µ_MAX_ = 0.21 ± 0.01 h^−1^), which is 56.8 % and 65.6 % of the corresponding rates presented by the indigenous strain UFMG-CM-Y259 and the industrial strain JP1, respectively (**Table 1**). Strain CEN.PK113-7D was also found to present the lowest specific substrate consumption rate (q_SMAX_ = −8.23 ± 0.37 mmol_GLCeq_ g_DM_^−1^ h^−1^), when compared to the values displayed by UFMG-CM-Y259 (q_SMAX_ = −12.84 ± 0.21 mmol_GLCeq_ g_DM_^−1^ h^−1^) and by JP1 (q_SMAX_= −12.29 ± 0.12 mmol_GLCeq_ g_DM_^−1^ h^−1^).

**Table 1.**
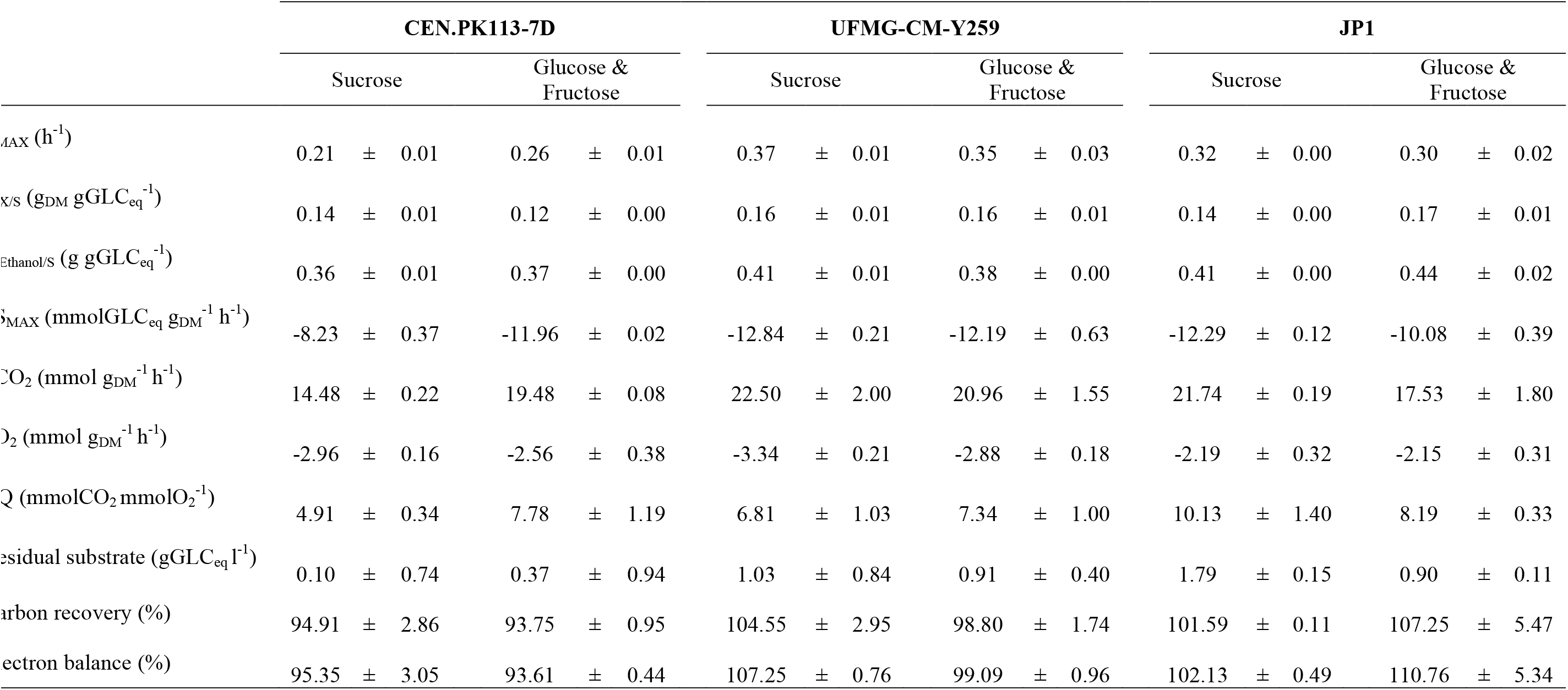
Relevant physiological parameters of *S. cerevisiae* CEN.PK113-7D, UFMG-CM-Y259, and JP1 during aerobic batch cultivations <u>with either sucrose or a glucose/fructose mixture as sole carbon and energy source</u>. All parameters were calculated for the exponential growth phase. Residual substrate refers to the remaining concentration of substrate at the end of the experiment. The data represent the mean of two experiments and the average deviation.

**Figure 2.**
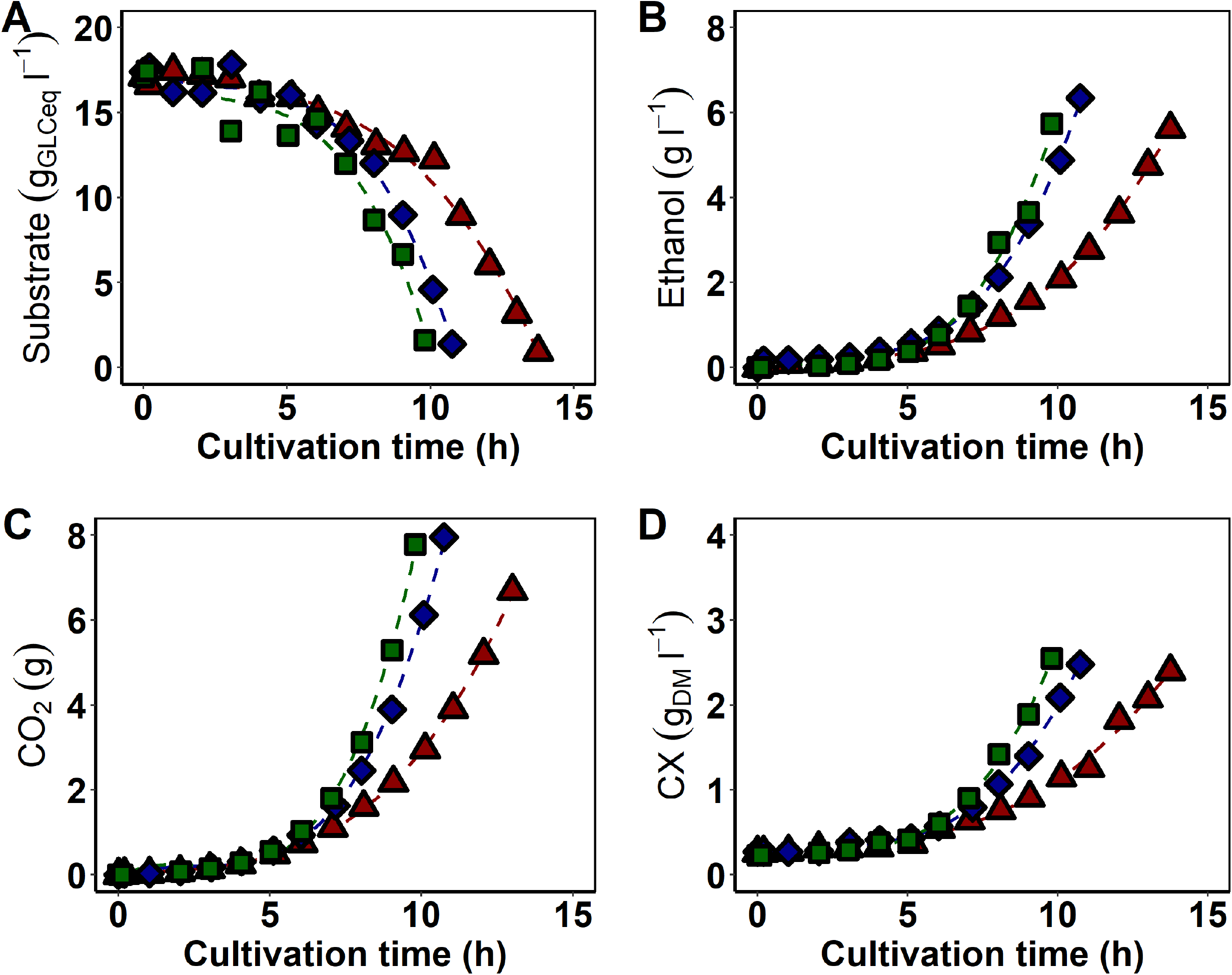
Substrate and metabolites concentrations/amounts during aerobic batch cultivation of *S. cerevisiae* CEN.PK113-7D (▴), JP1 (♦), and UFMG-CM-Y259 (▪) cells <u>with sucrose as sole carbon and energy source</u>. Substrate represents the sum of sucrose, glucose and fructose concentrations. Dashed lines represent trend lines. One representative dataset of duplicate independent experiments is shown.

**Figure 3.**
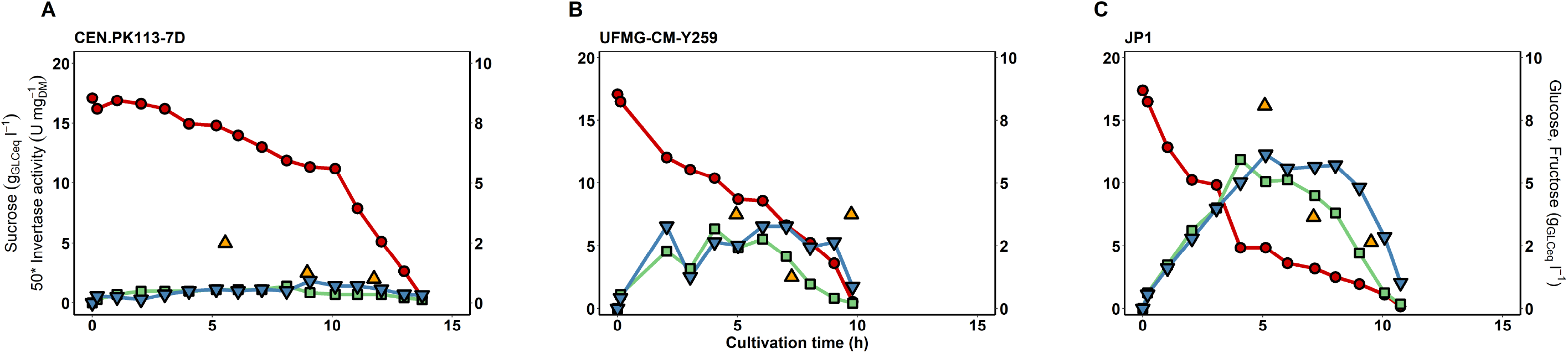
Sugar concentrations and periplasmic invertase activity during aerobic batch cultivation of *S. cerevisiae* CEN.PK113-7D (A), UFMG-CM-Y259 (B) and JP1 (C) cells on synthetic medium with <u>sucrose as sole carbon and energy source</u>. Sucrose (•); glucose (▪); fructose (*Closed ∇*); invertase (▴). Experiments were performed in duplicate. One representative dataset of duplicate independent experiments is shown.

Accordingly, the relative amount of substrate metabolized through the fermentative pathway was higher for UFMG-CM-Y259 and JP1 strains (~ 80%; **Supplementary Figure S1**) than that of CEN.PK113-7D (~ 70%; **Supplementary Figure S1**). This could also be deduced by the respiratory quotient (**Table 1**). In addition, allied to such observations, the ethanol yield on substrate was lower in the laboratory strain (Y_E/S_ = 0.36 ± 0.01 g g_GLCeq_^−1^), as compared to the indigenous (Y_E/S_ = 0.41 ± 0.01 g g_GLCeq_^−1^) and the industrial strain (Y_E/S_ = 0.41 ± 0.00 g g_GLCeq_^−1^).

One of the main traits of the yeast *Saccharomyces cerevisiae* is its capability to perform aerobic fermentation (De Deken 1966). In other words, even in the presence of oxygen, a higher relative amount of the substrate is subjected to fermentative catabolism rather than respiratory. A phenomenon that constrains the use of the respiratory pathway in yeast under circumstances of high sugar concentrations, as imposed in batch mode, is the so-called glucose (or carbon catabolite) repression. High concentrations of this sugar trigger a signaling cascade that will culminate in the repression of the transcription of genes encoding components of the electron transport chain and other respiratory proteins (Gancedo 1992; Santangelo 2006; Belinchón and Gancedo 2007a; Conrad et al. 2014). This strategy results in an energetically less efficient metabolism, as the ATP yield from fermentation is lower than that from respiration; for instance, in *S. cerevisiae* approximately 18 ATP are produced per mole of glucose via respiration against only 2 through fermentation (Verduyn et al. 1991; De Kok et al. 2012). A few hypotheses have been proposed to explain how cells overcome the lower ATP yield from fermentation (Pfeiffer and Morley 2014; Nilsson and Nielsen 2016; Niebel et al. 2019). The so-called rate/yield trade-off hypothesis (RYT), for instance, proposes that the cells accelerate growth to rise energy production rates (Pfeiffer and Morley 2014). Through another perspective, the use of fermentation allows for a higher ATP yield per protein mass as compared to respiration (Nilsson and Nielsen 2016). Our data somehow fit with these theories, in the sense that the lower growth rate of CEN.PK113-7D on sucrose, as compared to the other two strains, correlates with a lower relative amount of substrate channeled to fermentation in this laboratory strain.

The different ways in which the three yeast strains investigated here consume sucrose is likely to reflect the ecological niche of each one individually, at least partially. Strain JP1 was isolated from a sugarcane-based distillery where sugarcane juice — rather than molasses, which is not as sucrose-rich — is used to prepare the fermentation medium (da Silva Filho et al. 2005), meaning that it is adapted to an anaerobic environment, in which fermentative metabolism is the only option, and to excess sucrose. On the other hand, CEN.PK113-7D is commonly employed in fundamental laboratory research on physiology (van Dijken et al. 2000), where glucose is the major substrate. The long-term exposure of a microbe to a specific environment may alter its regulatory mechanisms, as well as cause adaptation, strategies used to cope with stress and to enhance fitness. In this sense, JP1 holds great advantage over the CEN.PK113-7D strain when sucrose is the carbon and energy source, due to its primary extensive contact with this sugar, which may have changed sucrose regulation in this lineage, or triggered mutations in genes encoding proteins that are essential to sucrose metabolism, resulting in an improved phenotype. We compared the *SUC2* sequences of JP1, UFMG-CM-Y259 and CEN.PK113-7D to the reference sequence of the S288c strain (Supplementary **Figure S6, Supplementary Table S4**). The protein sequence of the CEN.PK113-7D strain is identical to the one of the S288c strain. There are two amino acido changes shared by JP1 and UFMG-CM-Y259, namely N84H (Asn → His) and Q88E (Gln → Glu). In the JP1 strain alone, there are also two additional amino acid changes, with respect to the reference: A77T (Ala → Thr) and A409P (Ala → Pro). All other nucleotide differences do not lead to amino acid changes. We also analysed the 1,000 bp region upstream of the *SUC2* ORF (**Supplementary Table S5**) and verified that there are some differences when the JP1 and the UFMG-CM-Y259 sequences are compared to the reference. Whether the amino acid changes in the *SUC2* protein and/or any differences in gene expression due to nucleotide changes in the upstream region of *SUC2* in the JP1 and the UFMG-CM-Y259 strains are related to their faster growth phenotype on sucrose, when compared to the CEN.PK113-7D strain, remains to be experimentally tested.

Whether CEN.PK113-7D cells could evolve on sucrose and achieve a fitness comparable (or superior) to that of JP1 or UFMG-CM-Y259, remains to be explored. Previous laboratory evolution studies have demonstrated the potential of *S. cerevisiae* CEN.PK113-7D cells to improve its phenotype for maltose (Jansen et al. 2004), galactose (Hong et al. 2011) and mixed-substrate consumption (Papapetridis et al. 2018). Especially when sucrose was the substrate of choice, evolved engineered *S. cerevisiae* strains were shown to increase sucrose transport capacity (Basso et al. 2011; Marques et al. 2018). Concerning the UFMG-CM-Y259 strain, it was originally isolated from the barks of a tree that was brought by Europeans to Brazil, namely *Quercus rubra* (Northern Red Oak tree) (Beato et al. 2016). The characteristics of this niche are largely unknown. However, sucrose has been found in the barks of several other *Quercus* species from where *Saccharomyces* yeasts, including *S. cerevisiae*, have been isolated (Sampaio and Gonçalves 2008), suggesting that this disaccharide might also be present in Northern red oak tree’s bark. Also, another study showed that a yeast strain isolated from an oak tree performed better than Ethanol Red (a widespread industrial strain used mainly in fuel ethanol production from corn) under mimicked industrial conditions in the laboratory (Ruyters et al. 2015).

Taken together, these observations led us to believe that the historical background plays a key role on the enhanced growth and sucrose consumption displayed by JP1 and UFMG-CM-Y259, over CEN.PK113-7D cells.

### During cultivations of *S. cerevisiae* on sucrose, higher glucose and fructose accumulation seem to correlate with higher growth rates

The sugar consumption profiles during aerobic growth on sucrose of the three strains reveal that the accumulation of the released monosaccharides in the cultivation broth occurs to different extents (**Figure 3**). The highest concentrations, 6.0 g_GLCeq_.l^−1^ of glucose and 6.1 g_GLCeq_.l^−1^ of fructose, were observed for the industrial strain JP1. For UFMG-CM-Y259, the levels of extracellular glucose and fructose reached 3.2 g_GLCeq_.l^−1^ and 3.3 g_GLCeq_.l^−1^, respectively. On the other hand, the accumulation of hexoses was minimal with the laboratory strain CEN.PK113-7D and did not surpass 0.69 g_GLCeq_ l^−1^ and 0.91 g_GLCeq_ l^−1^ of glucose and fructose, respectively. Since this was the strain with the lowest maximum specific growth rate on sucrose (**Table 1**), we speculate that hexose accumulation during growth on this disaccharide contributes to faster growth. This, in turn, could be caused by the regulatory mechanisms related to the extracellular glucose concentration – such as the repression of the transcription of respiratory-related genes, which is known to have pleiotropic effects on yeast’s metabolism (Newcomb et al. 2003; Santangelo 2006; Belinchón and Gancedo 2007b; Kayikci and Nielsen 2015).

Moreover, as the accumulation of either glucose or fructose in the cultivation broth is a result of the difference between invertase activity and hexose transport rate, the observed different profiles could indicate that (1) hexose transport occurs at different rates in the different strains and/or (2) the enzyme invertase has different activity in each strain.

The existence of low- and high-affinity hexose transporter systems in the yeast *S. cerevisiae* has been well documented and the expression of these proteins depends on the levels of their substrates in the environment (Bisson et al. 1993; Ozcan and Johnston 1995; Kruckeberg 1996; Özcan and Johnston 1999). For low-affinity hexose transporters, such as Hxt1p and Hxt3p, the K_M_ for glucose ranges from 15 to 20 mM, while the K_M(glucose)_ of high-affinity hexose transporters, for instance Hxt2p and Hxt6p, is in the range of 1 to 2 mM (Özcan and Johnston 1999). Thus, one could assume that the cultivations on sucrose performed in this work triggered the expression of hexose transporters that differ from case to case, according to the concentration of glucose or fructose present in the broth. For instance, with the CEN.PK113-7D strain, for which the lowest levels of hexoses were observed (**Figure 3**), it is expected that high-affinity transporters are expressed. In a similar fashion, during cultivations with the UFMG-CM-Y259 and JP1 strains, low-affinity hexose transporters might be present. Transporter affinity and maximum velocity (V_max_) of these transport systems are inversely correlated in *S. cerevisiae* (Walsh et al. 1994). This means that a high-affinity transporter system displays lower V_max_ as compared to low-affinity, high-rate transporter systems. The hypothesis that hexose transport occurs at different rates in the three scenarios investigated here is, therefore, supported by these previously described properties of hexose transporters in *S. cerevisiae*.

### Invertase activity might constrain *S. cerevisiae* CEN.**PK113-7D’s growth rate on sucrose**

To investigate whether invertase activity contributes to the different hexose accumulation levels in the broth, we determined the specific activity of the periplasmic form of the enzyme in each of the three strains investigated here, during cultivations on sucrose, in the beginning, mid, and late exponential phase of growth.

In all samples, the periplasmic invertase of *S. cerevisiae* CEN.PK113-7D displays lower biomass specific activity, when compared to the UFMG-CM-Y259 and JP1 strains (**Figure 3**). The biomass specific periplasmic invertase activity achieved with the laboratory strain ranged from 0.04 ± 0.00 to 0.09 ± 0.00 U mg_DM_^−1^. In a previous study, Herwig and colleagues (Herwig et al. 2001) reported biomass specific total invertase activity within 0.2 to 1.0 U mg_DM_^−1^, approximately, for the *S. cerevisiae* CEN.PK113-7D strain cultivated in aerobic batch bioreactors at 30°C and pH 5.0, with sucrose as sole carbon and energy source. For this particular strain, it is known that the cytoplasmic invertase displays much lower activity compared to that of the periplasmic invertase (Basso et al. 2011; Marques et al. 2017), therefore the specific activity of the periplasmic form of this enzyme can be assumed to correspond roughly to total invertase activity. Hence, a remarkable difference (one order of magnitude) can be observed between the above mentioned study and our data, obtained under similar conditions. This was accompanied by much higher (approximately 10 times) levels of glucose and fructose in the broth observed by those authors, which is probably a consequence of the higher invertase activity achieved in their study. It is worth noticing that while our pre-cultivation was carried out with sucrose as sole carbon and energy source, their pre-culture was grown on glucose. This shift in the substrate from glucose in the pre-cultivation to sucrose in the cultivation itself likely explains the higher invertase activity observed by Herwig and colleagues, and is presumably a consequence of the cells being exposed to a sudden need for invertase.

The inferior biomass specific invertase activity combined with the lower levels of hexose accumulation in the broth (**Figure 3**) and the lower specific sucrose consumption rate (**Table 1**), suggests that sucrose hydrolysis is a constraint for CEN.PK113-7D’s growth on this carbon source. The two forms of the enzyme invertase (E.C. 3.2.1.2.6) present in *S. cerevisiae*’s cells are translated from genes of the *SUC* family that contains a total of nine structural genes (*SUC1*-*SUC5, SUC7*-*SUC10*) located in distinct loci of several chromosomes (Carlson et al. 1981; Carlson and Botstein 1983; Naumov and Naumova 2010). The *SUC2* gene, located in the subtelomeric region of chromosome IX (Carlson and Botstein 1983; Naumov and Naumova 2010), is found in all strains of the *Saccharomyces cerevisiae* species, as well as in other yeasts from the same genus (Naumov and Naumova 2010). The observed differences in biomass specific invertase activity among the strains here analyzed could be due to the mutations in the coding sequence for the *SUC2* gene of the UFMG-CM-Y259 and JP1 strains, as discussed above. We searched for the presence of additional *SUC* genes in the genomes of the JP1 and the UFMG-CM-Y259 strains. The analysis performed (results not shown) indicated no evidence for the presence of any *SUC* gene besides *SUC2*, and we could only find evidence for the presence of a single *SUC2* copy in both JP1 and UFMG-CM-Y259.

### The enhanced glycolytic rate of *S. cerevisiae* JP1 on sucrose might be due to a combined mechanism of sucrose utilization

To investigate whether the mechanism of sucrose utilization is responsible for the distinct substrate uptake rates observed in the sucrose-experiments, we carried out, under identical operational conditions, a physiological analysis of the studied strains on an equimolar mixture of glucose and fructose (**Figure 4; Table 1; Supplementary Figures S1-S5, Supplementary Tables S1-S3**). We hypothesized that if sucrose was being utilized exclusively via periplasmic hydrolysis, no difference in sugar uptake would be observed between the sucrose and the glucose/fructose mixture growth conditions, as long as no limitations in the hydrolysis step occurred.

**Figure 4.**
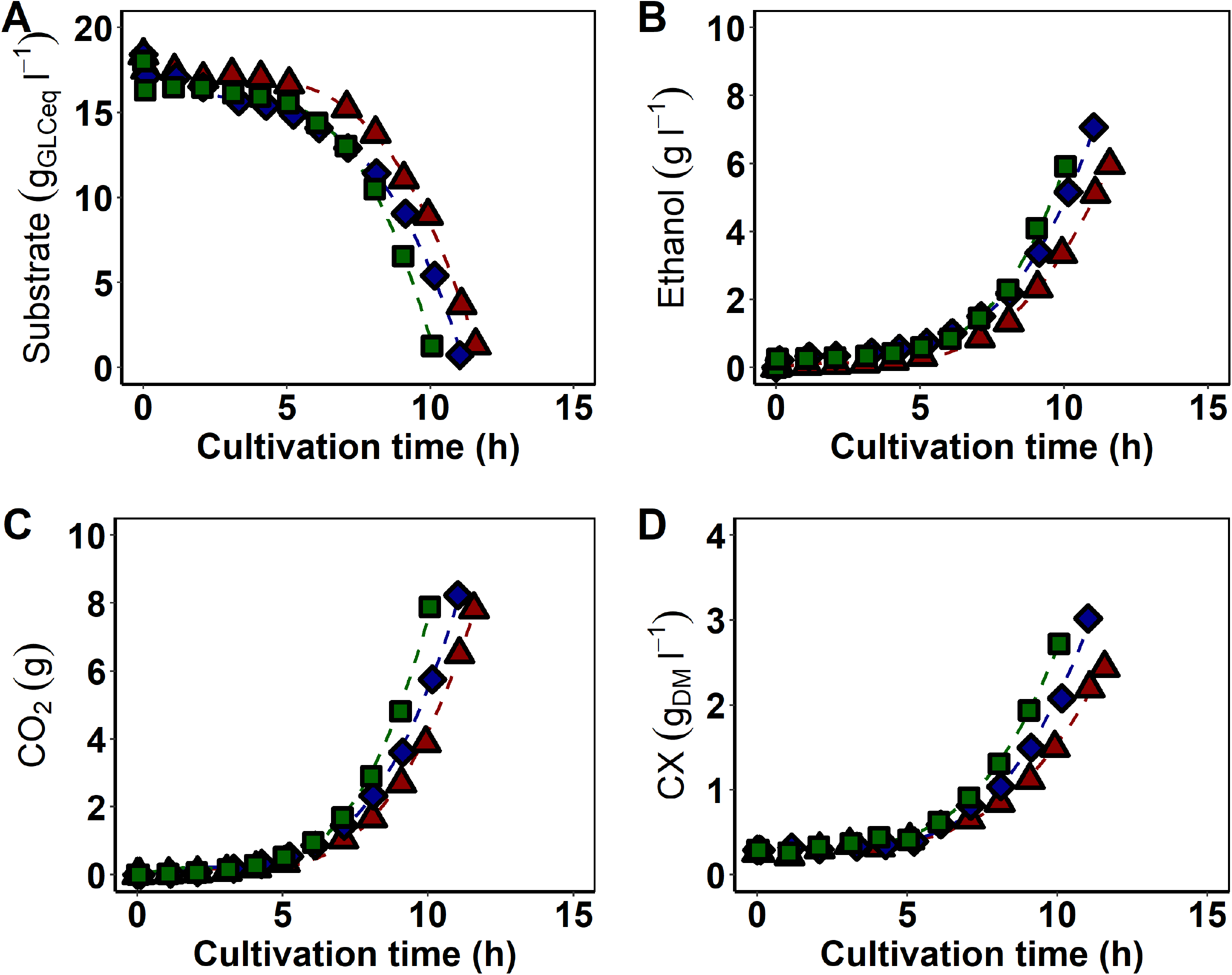
Substrate and metabolites concentrations/amounts during aerobic batch cultivation of *S. cerevisiae* CEN.PK113-7D (▴), JP1 (♦), and UFMG-CM-Y259 (▪) cells with an <u>equimolar mixture of glucose and fructose as sole carbon and energy source</u>. Substrate represent the sum of glucose and fructose concentrations. Dashed lines represent trend lines. One representative dataset of duplicate independent experiments is shown.

Furthermore, since the metabolism of sucrose in the yeast *S. cerevisiae* differs from that of its monomers (or a mixture of them) only in the step of disaccharide breakdown, when compared to growth on glucose and/or fructose, it can be speculated that active transport of sucrose accelerates the sugar uptake rate, and, consequently, the higher growth rate observed on the disaccharide. In fact, Barford and co-workers (Orlowski and Barford 1991; Barford et al. 1993) demonstrated that the superior growth of *S. cerevisiae* 248 UNSW 703100 — fully adapted to the culture medium for 20 - 250 generations — on sucrose, in comparison to a mixture of its monomers, was due to the direct uptake of sucrose molecules by actively growing yeast cells, which is faster than the passive transport of hexoses.

From our experiments, *S. cerevisiae* CEN.PK113-7D cells were more efficient in consuming the hexoses when they were provided directly (q_SMAX_sucrose_ = −8.23 ± 0.37 mmol_GLCeq_ g_DM_^−1^ h^−1^, q_SMAX_G+F_ = −11.96 ± 0.02 mmol_GLCeq_ g_DM_^−1^ h^−1^; **Table 1; Table S1**); the opposite was observed with the industrial strain, JP1 (q_SMAX_sucrose_ = −12.29 ± 0.12 mmol_GLCeq_ g_DM_^−1^ h^−1^, q_SMAX_G+F_ = −10.08 ± 0.39 mmol_GLCeq_ g_DM_^−1^ h^−1^; **Table 1; Table S3**), whereas the UFMG-CM-Y259 strain displayed equivalent sugar uptake rates for the sucrose experiment, as compared to the one with the glucose/fructose mixture (q_SMAX_sucrose_ = −12.84 ± 0.21 mmol_GLCeq_ g_DM_^−1^ h^−1^, q_SMAX_G+F_ = −12.19 ± 0.63 mmol_GLCeq_ g_DM_^−1^ h^−1^; **Table 1; Table S2**).

The observation of immediate glucose and fructose formation in the broth as well as the measured biomass specific periplasmic invertase activity indicate that extracellular hydrolysis of sucrose is a mechanism of sucrose utilization in all experiments carried out on sucrose alone. However, the hypothesis that sucrose is not being actively transported likely holds true for the yeast strains UFMG-CM-Y259 and CEN.PK113-7D. In the latter case, as discussed above, there is evidence for a growth limitation caused by insufficient invertase activity. Because sucrose-grown JP1 cells displayed enhanced glycolytic rates, when compared to cells cultivated on the glucose/fructose mixture — which is evidenced both by the specific rates of substrate consumption and by the specific rates of products formation (**Table 1; Table S3**) —, we believe that a combined mechanism of sucrose utilization is likely to take place when JP1 cells grow on sucrose alone. In other words, both periplasmic hydrolysis by invertase and active transport of sucrose occur in parallel (**Figure 1**).

This assumption is supported by a previous work from Barford et al. (Barford et al. 1992), in which a combination of direct sucrose uptake and extracellular hydrolysis with subsequent transport of its monomers was proposed to explain the higher glycolytic rate of the evolved *S. cerevisiae* 248 UNSW 703100 strain when grown on sucrose, as compared to an equimolar mixture of the hexoses. Moreover, recently, Prado and co-workers (Prado et al. 2020) argued that a mixed mode of sucrose utilization by the *S. cerevisiae* strain LBGA-01 was responsible for its improved performance at high temperature.

Since the reference *S. cerevisiae* S288c strain does not have a functional *AGT1* allele, which is incorrectly annotated as a synonym of *MAL11* in the yeastgenome.org database (Trichez et al. 2019), we used as a reference the *AGT1* sequence of the industrial *S. cerevisiae* CAT-1 strain (Babrzadeh et al. 2012). The JP1 and the UFMG-CM-Y259 strains presente several (> 20) amino acid changes, with respect to the sequence of the CAT-1 strain (**Supplementary Table S6 and Figure S7**). Again, these differences remain to be explored experimentally to see whether *AGT1* in the two strains investigated here encodes an improved sucrose transporter.

### Sucrose-grown *S. cerevisiae* UFMG-CM-Y259 cells display higher maximum specific growth rate than glucose-grown cells

A previous study from our group (Beato et al. 2016) revealed the capacity of some *S. cerevisiae* strains, including UFMG-CM-Y259, to grow faster on sucrose than on glucose through experiments carried out using microtiter plates as a cultivation system and optical density measurements for assessing cell concentration. As the calculation of the maximum specific growth rate can lead to different values depending on e.g. the cultivation system and the cell concentration measurements used (Stevenson et al. 2016), we sought to investigate whether such observations would hold in a well-controlled cultivation system, i.e. a bioreactor combined with direct cell concentration measurements via dry mass determinations. We performed cultivations of the three *S. cerevisiae* strains on glucose as the sole carbon and energy source (**Figure 5; Table 2; Supplementary Figures S1-S5, Supplementary Tables S1-S3**), otherwise under identical conditions as the cultivations hitherto discussed. The UFMG-CM-Y259 strain displayed higher maximum specific growth rate on sucrose (µ_MAX_ = 0.37 ± 0.01 h^−1^; **Table 1; Table S2**) than on glucose (µ_MAX_ = 0.29 ± 0.00 h^−1^; **Table 2; Table S2**), which corroborates the aforementioned study. Results for the JP1 strain from both studies are also in agreement, in the sense that the strain presented a slightly higher µ_MAX_ on sucrose than on glucose, although this difference was not statistically significant in the previous study (at the 95% confidence level, triplicate experiments in (Beato et al. 2016)). In the present study, we did not perform statistical analysis, since we only performed duplicate experiments in bioreactors, but µ_MAX_ on sucrose and on glucose for JP1 was 0.32 and 0.28 h^−1^, respectively. For the CEN.PK113-7D strain, while the results from the previous study indicated no statistical difference between µ_MAX_ on sucrose and on glucose, here the value on sucrose (0.21 h^−1^) was much lower than on glucose (0.31 h^−1^). We speculate that this could be due to the fact that in the previous study (Beato et al. 2016), absorbance measurements, instead of the dry cell mass measurements employed here, were used to calculate µ_MAX_. Since the CEN.PK113-7D strain is haploid, and haploid cells are typically smaller than diploid ones, the wavelength used in spectrophotometric analysis (typically around 600 nm) is closer to the size of the cells, causing bigger differences between direct cell concentration and turbidity measurements (Stevenson et al. 2016).

**Table 2.** Relevant physiological parameters of *S. cerevisiae* CEN.PK113-7D, UFMG-CM-Y259, and JP1 during aerobic batch cultivations <u>with either glucose or fructose as sole carbon and energy source</u>. All parameters were calculated for the exponential growth phase. Residual substrate refers to the remaining substrate concentration at the end of the experiment. The data represent the mean of two experiments and the average deviation.

|  | CEN.PK113-7D |  |  |  | UFMG-CM-Y259 |  |  |  | JP1 |  |  |  |
| --- | --- | --- | --- | --- | --- | --- | --- | --- | --- | --- | --- | --- |
|  | Glucose |  | Fructose |  | Glucose |  | Fructose |  | Glucose |  | Fructose |  |
| $\mu_{MAX}$ (h<sup>-1</sup>) | 0.31 | ± 0.01 | 0.32 | ± 0.01 | 0.29 | ± 0.00 | 0.36 | ± 0.02 | 0.28 | ± 0.02 | 0.32 | ± 0.02 |
| $\gamma_{X/S}$ (g<sub>DM</sub> gGLC<sub>eq</sub><sup>-1</sup>) | 0.13 | ± 0.01 | 0.13 | ± 0.00 | 0.14 | ± 0.00 | 0.16 | ± 0.00 | 0.13 | ± 0.01 | 0.15 | ± 0.01 |
| $\gamma_{Ethanol/S}$ (g gGLC<sub>eq</sub><sup>-1</sup>) | 0.41 | ± 0.01 | 0.38 | ± 0.00 | 0.40 | ± 0.03 | 0.37 | ± 0.00 | 0.42 | ± 0.00 | 0.38 | ± 0.00 |
| $S_{MAX}$ (mmolGLC<sub>eq</sub> g<sub>DM</sub><sup>-1</sup> h<sup>-1</sup>) | -13.22 | ± 0.23 | -13.46 | ± 0.51 | -11.96 | ± 0.43 | -12.12 | ± 0.84 | -11.67 | ± 0.45 | -12.24 | ± 0.07 |
| $\gamma_{CO_2}$ (mmol g<sub>DM</sub><sup>-1</sup> h<sup>-1</sup>) | 23.57 | ± 0.22 | 23.48 | ± 0.34 | 20.77 | ± 0.50 | 20.44 | ± 1.39 | 19.75 | ± 0.31 | 20.89 | ± 0.05 |
| $\gamma_{O_2}$ (mmol g<sub>DM</sub><sup>-1</sup> h<sup>-1</sup>) | -2.45 | ± 0.14 | -2.84 | ± 0.11 | -2.63 | ± 0.31 | -3.03 | ± 0.12 | -2.12 | ± 0.09 | -2.43 | ± 0.02 |
| $\gamma_Q$ (mmolCO<sub>2</sub> mmolO<sub>2</sub><sup>-1</sup>) | 9.66 | ± 0.66 | 8.28 | ± 0.19 | 8.01 | ± 1.12 | 6.73 | ± 0.20 | 9.31 | ± 0.25 | 8.60 | ± 0.04 |
| Residual substrate (gGLC<sub>eq</sub>.l<sup>-1</sup>) | 0.15 | ± 0.12 | 2.72 | ± 1.35 | 0.03 | ± 0.03 | 2.28 | ± 0.19 | 0.73 | ± 0.08 | 1.34 | ± 0.10 |
| Carbon recovery (%) | 100.73 | ± 0.07 | 97.57 | ± 0.66 | 103.25 | ± 6.11 | 99.04 | ± 0.94 | 101.44 | ± 1.35 | 97.12 | ± 0.51 |
| Electron balance (%) | 101.14 | ± 0.70 | 97.41 | ± 0.01 | 99.05 | ± 2.05 | 100.00 | ± 0.96 | 104.06 | ± 0.53 | 97.25 | ± 0.66 |

**Figure 5.**
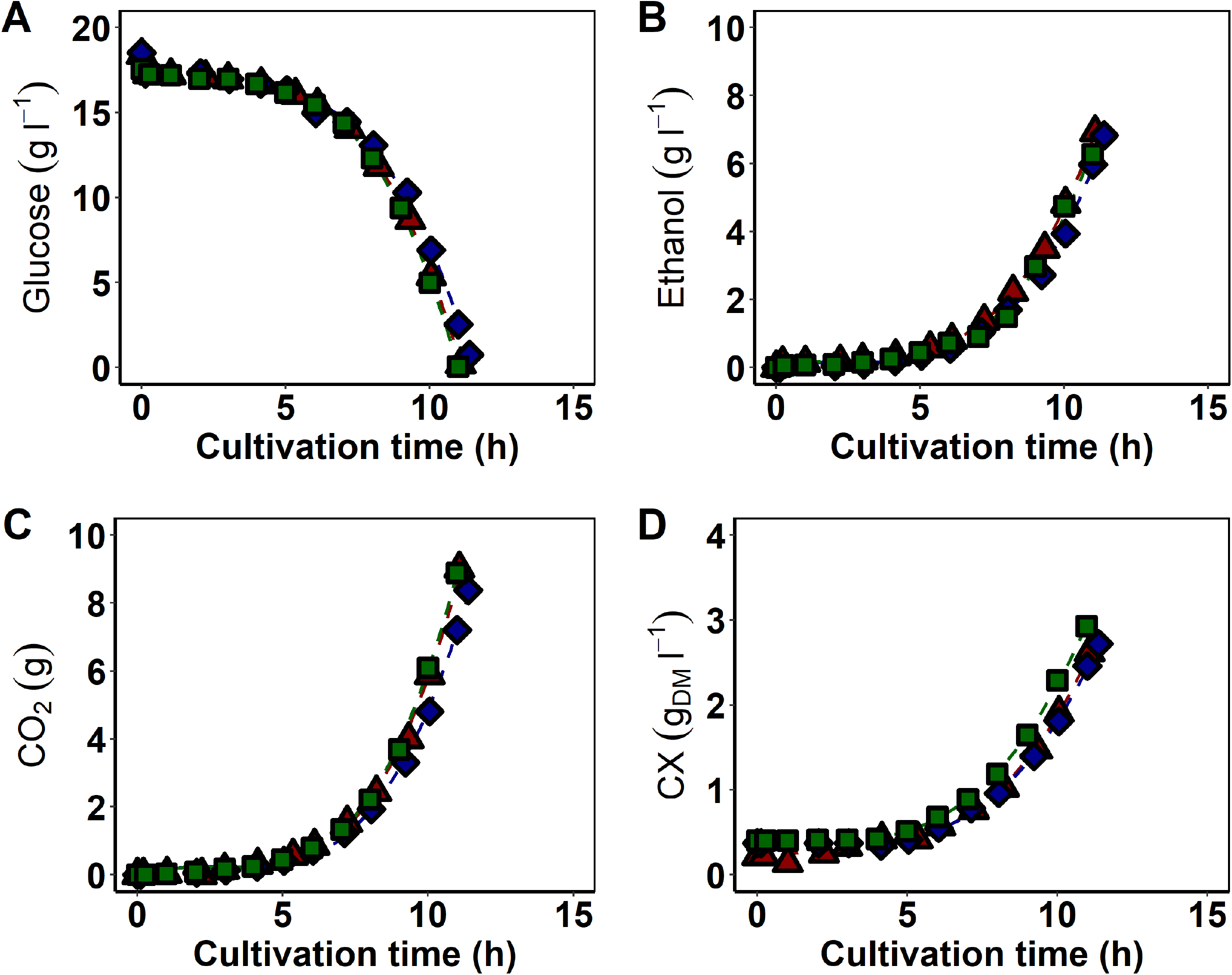
Substrate and metabolites concentrations/amounts during aerobic batch cultivation of *S. cerevisiae* CEN.PK113-7D (▴), JP1 (♦), and UFMG-CM-Y259 (▪) cells with <u>glucose as sole carbon and energy source</u>. Dashed lines represent trend lines. One representative dataset of duplicate independent experiments is shown.

In principle, the maximum specific growth rate on sucrose is not expected to exceed that on glucose because the yeast *S. cerevisiae* is subjected to glucose repression (Gancedo 1998), meaning that any carbon and energy source different from glucose eventually present in the medium will have their consumption delayed, as long as glucose is available. This preference mechanism suggests that the microbe will display a higher specific growth rate on glucose than on any other carbon and energy source. As previously discussed in this work, direct sucrose uptake could explain enhanced growth rates on this disaccharide over related carbon sources. Alongside the superior growth rate, we observed a slightly higher substrate uptake rate during sucrose cultivation (q_SMAX_sucrose_ = −12.84 ± 0.21 mmol_GLCeq_ g_DM_^−1^ h^−1^; **Table 1; Table S2**), as compared to growth on glucose alone (q_SMAX_glucose_ = −11.96 ± 0.43 mmol_GLCeq_ g_DM_^−1^ h^−1^; **Table 2; Table S2**) for the yeast UFMG-CM-Y259. The remaining physiological parameters were equivalent in both growth conditions.

Through another perspective, glucose and sucrose differently impact signaling cascades in the cells, such as the one leading to the regulation of the protein kinase A (PKA) activity. A theory related to the PKA signaling cascade has also been proposed to explain high specific growth rates of *S. cerevisiae* on sucrose (Badotti et al. 2008; Marques et al. 2016). This cascade is activated when the Gpr1-Gpa2 coupled receptor senses either glucose or sucrose, with the affinity of Gpr1p being higher for the latter sugar (Lemaire et al. 2004). Once activated, the PKA protein can exert all its function, including regulating the synthesis and degradation of storage carbohydrates (Wingender-Drissen and Becker 1983; WERA et al. 1999; François et al. 2012), the metabolic flux through the glycolytic pathway (Dihazi et al. 2003; Portela et al. 2006), and gluconeogenesis (Mazón et al. 1982).

To this point, the mechanisms behind faster growth on sucrose over glucose remain an open question. Further investigation, for instance by means of systems biology approaches, is needed to elucidate sucrose regulation in the yeast *S. cerevisiae*.

### *S. cerevisiae* UFMG-CM-Y259 and JP1 display similar physiology during growth on sucrose or on fructose

At last, we performed aerobic batch cultivations with fructose as the sole carbon and energy source (**Figure 6; Table 2; Supplementary Figures S1-S5, Supplementary Tables S1-S3**), again otherwise under identical conditions, when compared to all other cultivations described in this work. The physiological parameters were compared with those obtained from sucrose cultivations. The maximum specific growth rate between cultivations on the monosaccharide or on the disaccharide differed only for the CEN.PK113-7D strain, that grew faster when cultivated on the hexose (µ_max_Fructose_ = 0.32 ± 0.01 h^−1^ and µ_max_Sucrose_ = 0.21 ± 0.01 h^−1^). The industrial (JP1) and the indigenous (UFMG-CM-Y259) strains’ physiologies were equivalent when the carbon and energy sources sucrose or fructose are compared. Interestingly, the UFMG-CM-Y259 strain’s physiology on fructose was more similar to that on sucrose than to the performance on glucose, contrasting the observations for the CEN.PK113-7D strain. This suggests that even the regulatory mechanisms triggered by hexoses in the yeast *S. cerevisiae* are strain-dependent.

**Figure 6.**
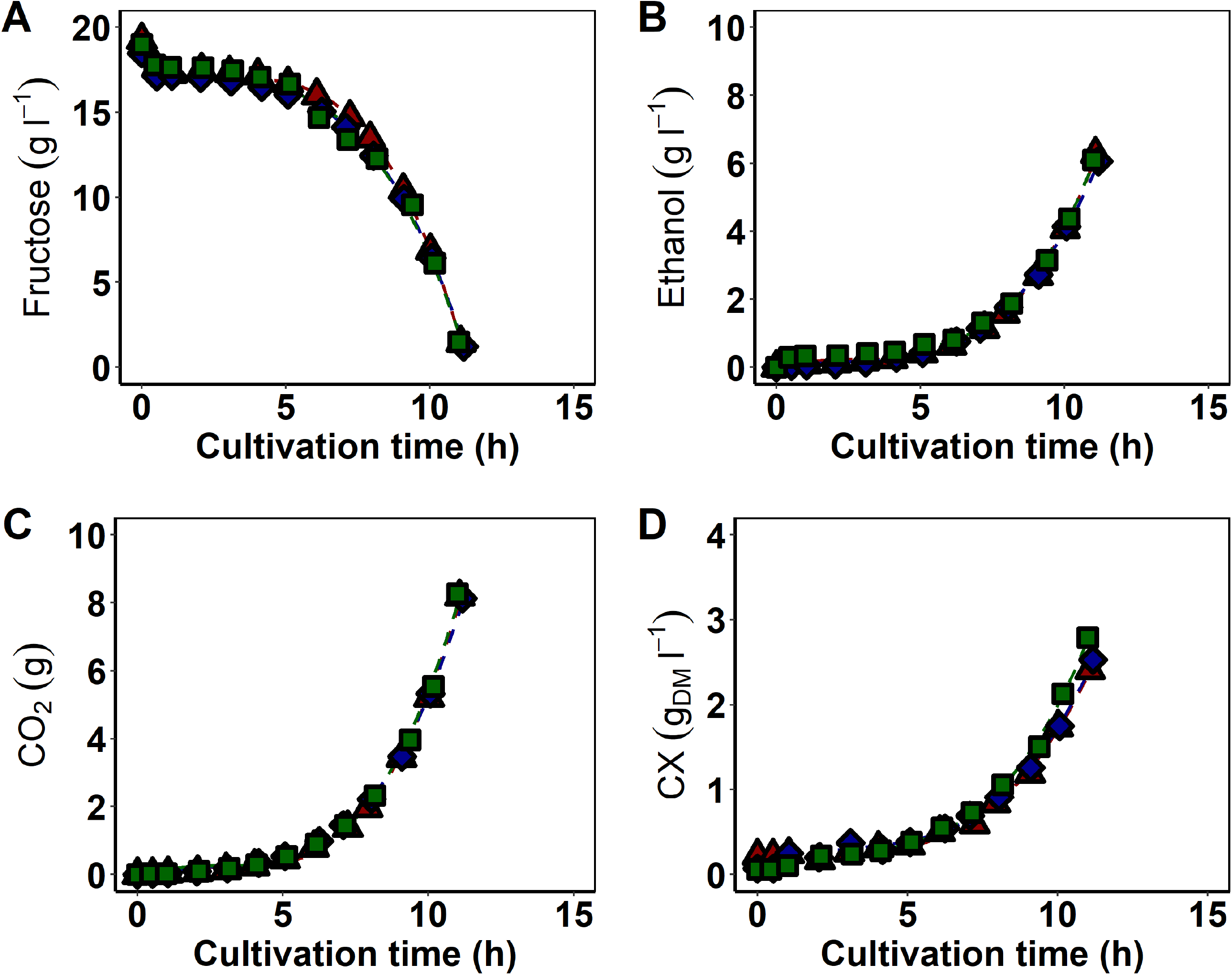
Substrate and metabolites concentrations/amounts during aerobic batch cultivation of *S. cerevisiae* CEN.PK113-7D (▴), JP1 (♦), and UFMG-CM-Y259 (▪) cells with <u>fructose as sole carbon and energy source</u>. Dashed lines represent trend lines. One representative dataset of duplicate independent experiments is shown.

Considering only glucose and fructose, transport has been pinpointed as a critical step for the different behaviors observed in the metabolism of such hexoses in *S. cerevisiae* (Luyten et al. 2002; Berthels et al. 2004). Among all the 20 proteins that constitute the group of hexose transporters and glucose sensors present in this microbe (Kruckeberg 1996; Özcan and Johnston 1999), Hxt1p to Hxt7p are the main ones in the context of glucose and/or fructose catabolism (Kruckeberg 1996; Luyten et al. 2002; Verwaal et al. 2002; Guillaume et al. 2007; Karpel et al. 2008). Expression of these proteins is strain dependent (Verwaal et al. 2002; PEREZ et al. 2005) and their substrate affinities vary from one transporter to the other (Ozcan and Johnston 1995). Enhanced fructose fermentation capacity has been demonstrated to be associated to the Hxt3p transporter (Guillaume et al. 2007). In the latter study (Guillaume et al. 2007), the molecular basis behind a higher fructose utilization capacity displayed by the commercial wine *S. cerevisiae* Fermichamp was revealed, in comparison to that of standard *S. cerevisiae* wine strains, namely the mutations T200A and G415N in the *HXT3* DNA sequence were responsible for such phenotype. Our sequence analysis of the *HXT3* open reading frame (ORF) did not show these same amino acid changes for any of the strains studied here (**Supplementary Figure S8, Supplementary Tables S7**). However, the T50A and V428C non-conservative amino acid substitutions were found in the *HXT3* ORF of the UFMG-CM-Y259 strain, and the K320Q substitution in this ORF in JP1. These could be leads to be followed to verify whether these *HXT3* alleles are capable of transporting fructose with different affinities and/or capacities.

Besides the transport step, hexose phosphorylation could also contribute to the different growth physiologies exhibited by the UFMG-CM-Y259 strain on the monosaccharides, since it is the first step in yeast glycolysis. *In vitro* measurements have shown that the proteins hexokinase 1 and 2 target both monosaccharides (glucose or fructose), with lower affinity and higher relative maximum velocity of reaction (V_max_) for fructose (Lobo and Maitra 1977). Glucokinase, on the other hand, is insensitive to fructose (Lobo and Maitra 1977). Among these isozymes, hexokinase 2 is the major kinase in this glycolytic step, as its absence dramatically changes *S. cerevisiae*’s physiology (Diderich et al. 2001). It is noteworthy that while the conversion of fructose into fructose-6-phosphate takes only one step, to convert glucose into the same metabolite two steps are required (phosphorylation and isomerization).

## ACKNOWLEDGEMENTS

We would like to acknowledge Fundação de Amparo à Pesquisa do Estado de São Paulo (FAPESP, São Paulo, Brazil) for funding and for scholarships to CISR (grant numbers 2016/07285-9, 2017/08464-7, 2017/18206-5). This work was carried out as part of a Dual Degree PhD project under the agreement between UNICAMP and Delft University of Technology.

## Notes

### Competing Interest Statement

The authors have declared no competing interest.

