## Supplementary for "Aerobic growth physiology of *Saccharomyces cerevisiae* on sucrose is strain-dependent"

**Supplementary Material (Figures and Tables)**

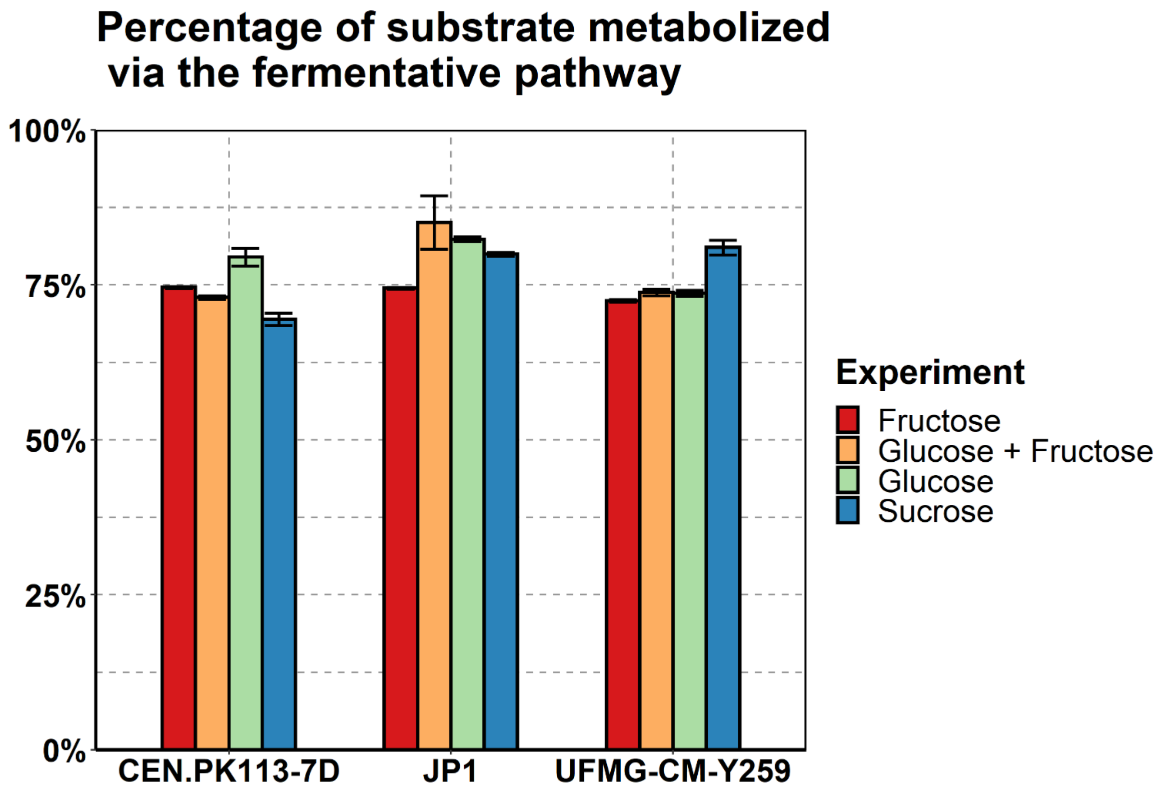


**Figure S1.** Percentage of substrate metabolized via the fermentative pathway by *S. cerevisiae* CEN.PK113-7D, UFMG-CM-Y259, and JP1 during aerobic growth on sucrose as sole carbon and energy source. Error bars represent the average deviation of the values obtained from duplicate experiments.

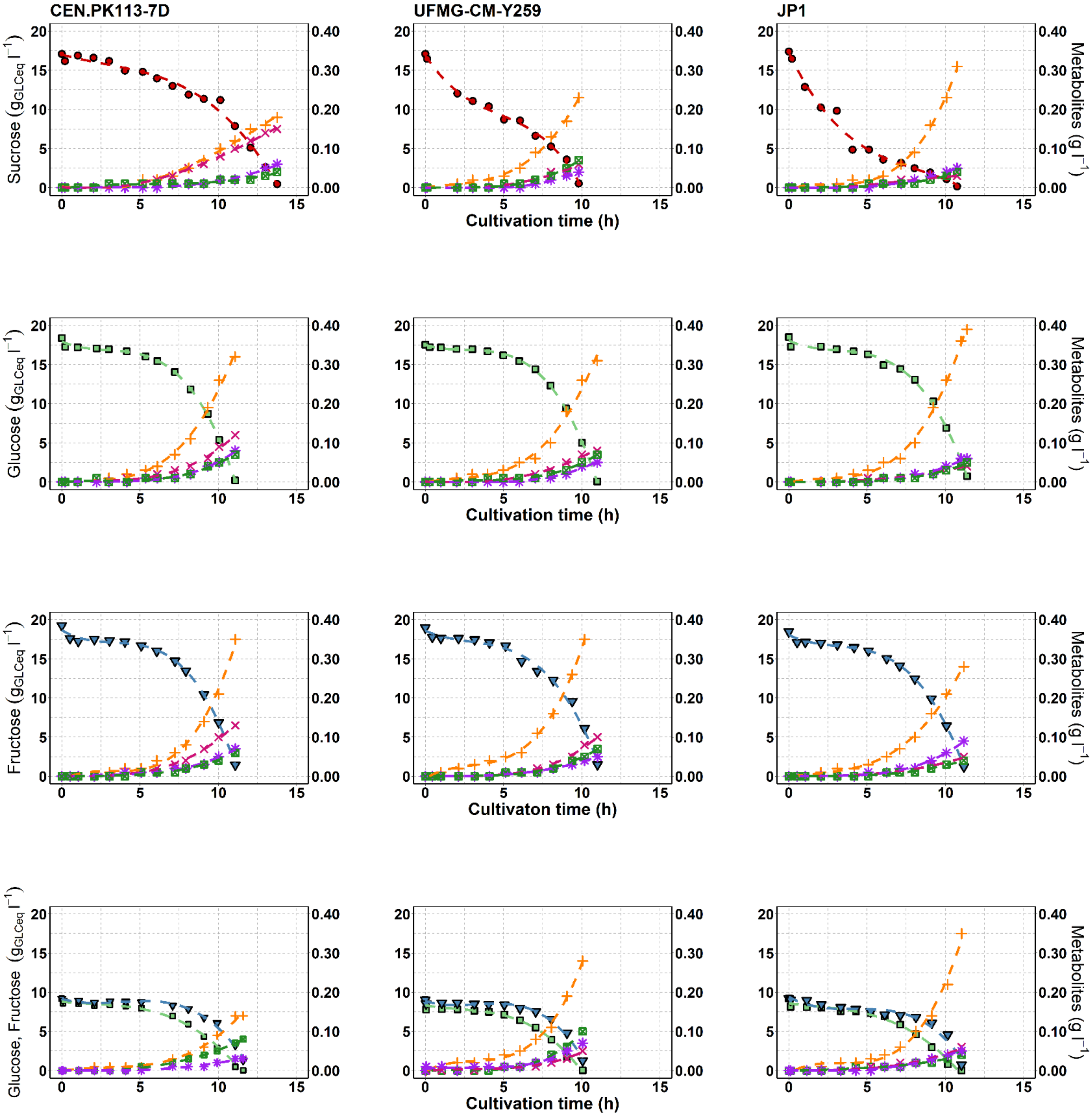


**Figure S2.** Substrate and metabolites concentrations during aerobic batch cultivation of S. cerevisiae CEN.PK113-7D, UFMG-CM-Y259 and JP1 with either sucrose, glucose, fructose or an equimolar mixture of glucose and fructose as sole carbon and energy source. Sucrose (●); Glucose

(■), Fructose ($closed\ \nabla$); Glycerol (+); Acetate (X); Succinate (*); Lactate (crossed □). Dashed lines represent trend lines. Experiments were performed in duplicate. Data shown in the plots are from a single growth experiment.

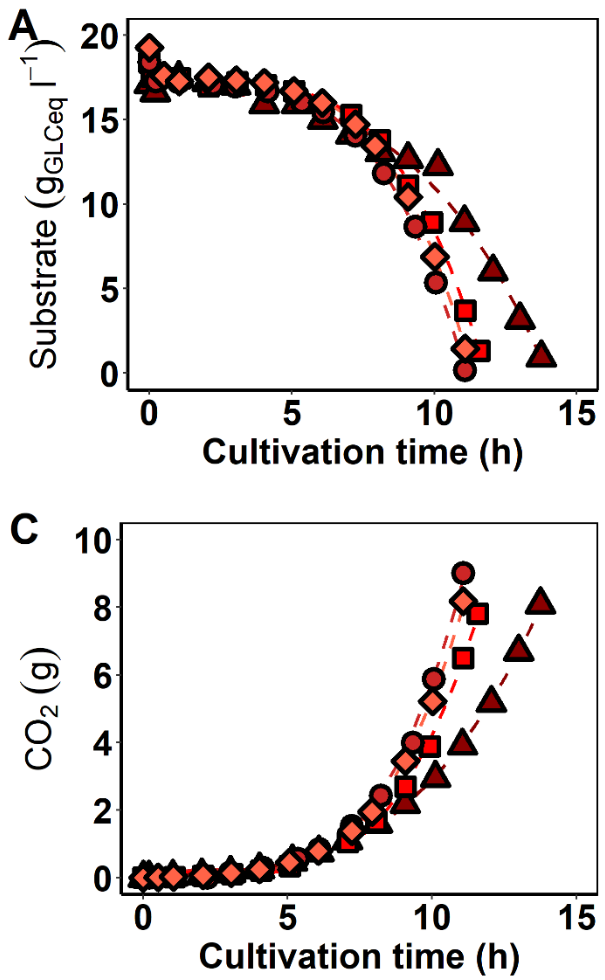


**Figure S3.** Substrate and metabolites concentrations/amounts during aerobic batch cultivation of *S. cerevisiae* CEN.PK113-7D with sucrose (▲), an equimolar mixture of glucose and fructose (■), glucose (●), or fructose (♦) as sole carbon and energy source. For the glucose + fructose experiment, substrate represents the sum of glucose and fructose concentrations. Dashed lines represent trend lines. Experiments were performed in duplicate. Data shown in the plots are from a single growth experiment.

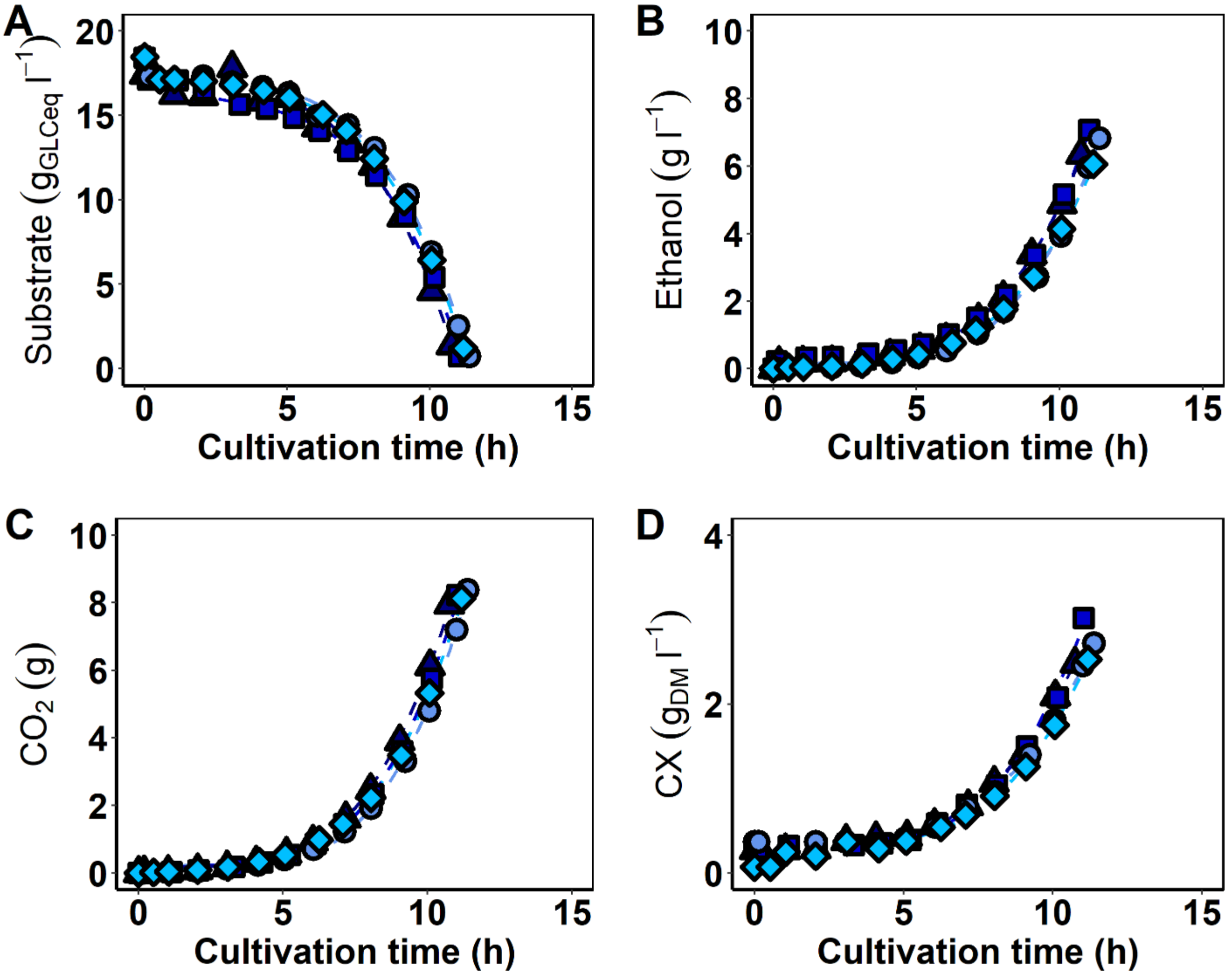


**Figure S4.** Substrate and metabolites concentrations/amounts during aerobic batch cultivation of *S. cerevisiae* JP1 with sucrose (▲), an equimolar mixture of glucose and fructose (■), glucose (●), or fructose (♦) as sole carbon and energy source. For the glucose + fructose experiment, substrate represents the sum of glucose and fructose concentrations. Dashed lines represent trend lines. Experiments were performed in duplicate. Data shown in the plots are from a single growth experiment.

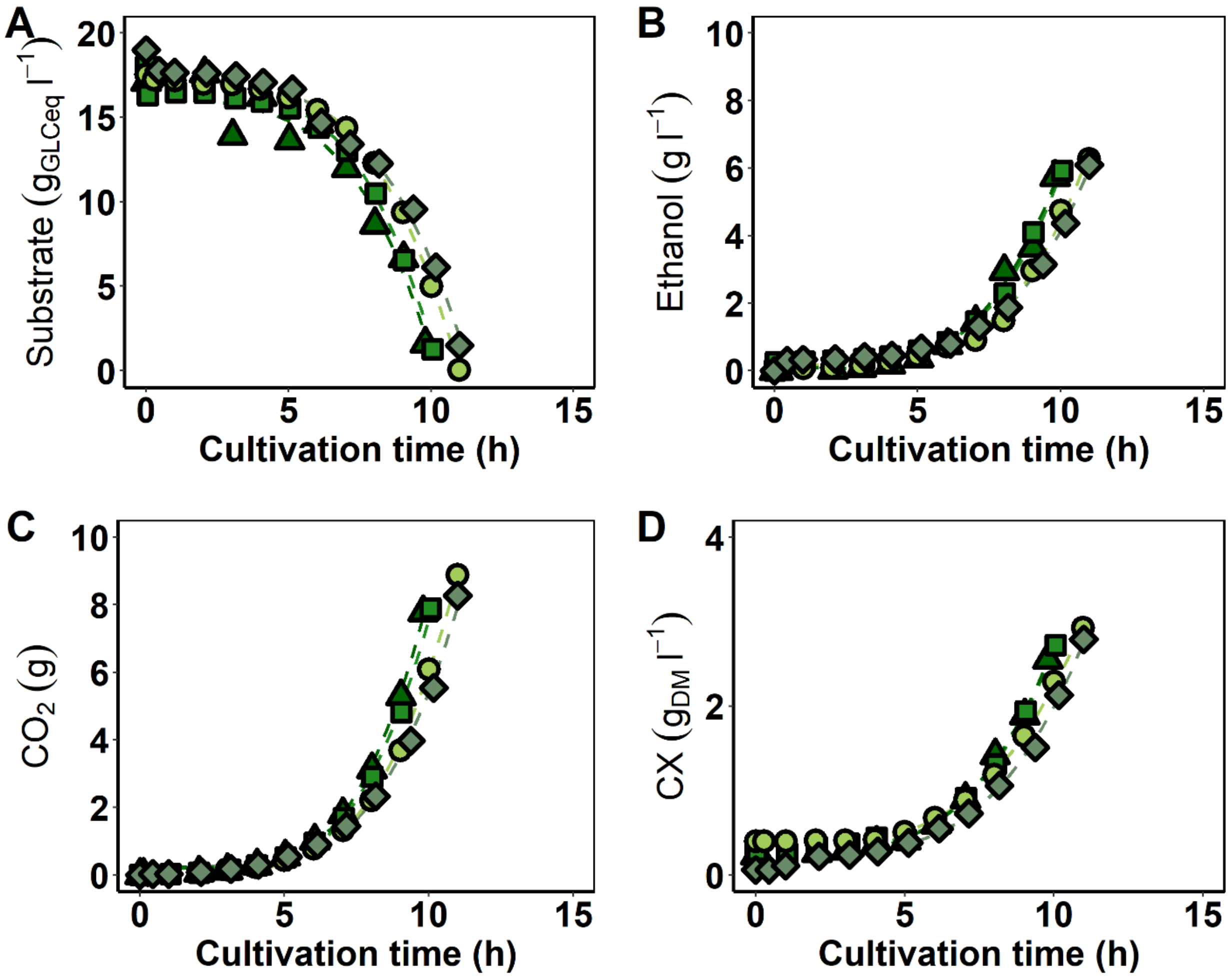


**Figure S5.** Substrate and metabolites concentrations/amounts during aerobic batch cultivation of *S. cerevisiae* UFMG-CM-Y259 with sucrose (▲), an equimolar mixture of glucose and fructose (■), glucose (●), or fructose (♦) as sole carbon and energy source. For the glucose + fructose experiment, substrate represents the sum of glucose and fructose concentrations. Dashed lines represent trend lines. Experiments were performed in duplicate. Data shown in the plots are from a single growth experiment.

**Table S1.** Physiological parameters of *S. cerevisiae* CEN.PK113-7D during aerobic batch cultivations with either glucose, sucrose, an equimolar mixture of glucose and fructose, or fructose as sole carbon and energy source. All parameters were calculated for the exponential growth phase. The data represent the mean of two experiments and the average deviation.

| | **Strain: CEN.PK113-7D** | | | |
|---|---|---|---|---|
| | Glucose | Sucrose | Glucose & Fructose | Fructose |
| $\mu_{MAX}$ ($h^{-1}$) | 0.31 ± 0.01 | 0.21 ± 0.01 | 0.26 ± 0.01 | 0.32 ± 0.01 |
| $Y_{X/S}$ ($g_{DM}$ $gGLC_{eq}^{-1}$) | 0.13 ± 0.01 | 0.14 ± 0.01 | 0.12 ± 0.00 | 0.13 ± 0.00 |
| $Y_{Ethanol/S}$ (g $gGLC_{eq}^{-1}$) | 0.41 ± 0.01 | 0.36 ± 0.01 | 0.37 ± 0.00 | 0.38 ± 0.00 |
| $Y_{CO2/S}$ (g $gGLC_{eq}^{-1}$) | 0.44 ± 0.00 | 0.43 ± 0.01 | 0.40 ± 0.00 | 0.43 ± 0.01 |
| $Y_{glycerol/S}$ (g $gGLC_{eq}^{-1}$) | 0.02 ± 0.00 | 0.01 ± 0.00 | 0.02 ± 0.01 | 0.03 ± 0.00 |
| $Y_{acetate/S}$ (g $gGLC_{eq}^{-1}$) | 0.01 ± 0.00 | 0.01 ± 0.00 | 0.01 ± 0.00 | 0.01 ± 0.00 |
| $Y_{succinate/S}$ (g $gGLC_{eq}^{-1}$) | 0.00 ± 0.00 | 0.00 ± 0.00 | 0.00 ± 0.00 | 0.00 ± 0.00 |
| $Y_{lactate/S}$ (g $gGLC_{eq}^{-1}$) | 0.00 ± 0.00 | 0.00 ± 0.00 | 0.00 ± 0.00 | 0.00 ± 0.00 |
| $qS_{MAX}$ ($mmolGLC_{eq}$ $g_{DM}^{-1}$ $h^{-1}$) | -13.22 ± 0.23 | -8.23 ± 0.37 | -11.96 ± 0.02 | -13.46 ± 0.51 |
| $qCO_2$ (mmol $g_{DM}^{-1}$ $h^{-1}$) | 23.57 ± 0.22 | 14.48 ± 0.22 | 19.48 ± 0.08 | 23.48 ± 0.34 |
| $qO_2$ (mmol $g_{DM}^{-1}$ $h^{-1}$) | -2.45 ± 0.14 | -2.96 ± 0.16 | -2.56 ± 0.38 | -2.84 ± 0.11 |
| RQ ($mmolCO_2$ $mmolO_2^{-1}$) | 9.66 ± 0.66 | 4.91 ± 0.34 | 7.78 ± 1.19 | 8.28 ± 0.19 |
| $q_{Ethanol}$ (mmol $g_{DM}^{-1}$ $h^{-1}$) | 21.03 ± 0.74 | 11.43 ± 0.34 | 17.46 ± 0.09 | 20.08 ± 0.81 |
| $q_{glycerol}$ (mmol $g_{DM}^{-1}$ $h^{-1}$) | 0.53 ± 0.02 | 0.19 ± 0.01 | 0.37 ± 0.12 | 0.72 ± 0.01 |
| $q_{acetate}$ (mmol $g_{DM}^{-1}$ $h^{-1}$) | 0.21 ± 0.05 | 0.32 ± 0.06 | 0.28 ± 0.12 | 0.26 ± 0.03 |
| $q_{succinate}$ (mmol $g_{DM}^{-1}$ $h^{-1}$) | 0.08 ± 0.00 | 0.04 ± 0.01 | 0.05 ± 0.02 | 0.07 ± 0.02 |
| $q_{lactate}$ (mmol $g_{DM}^{-1}$ $h^{-1}$) | 0.13 ± 0.06 | 0.02 ± 0.02 | 0.05 ± 0.00 | 0.08 ± 0.01 |
| Residual substrate ($gGLC_{eq}$ $l^{-1}$) | 0.15 ± 0.12 | 0.10 ± 0.74 | 0.37 ± 0.94 | 2.72 ± 1.35 |
| Carbon recovery (%) | 100.73 ± 0.07 | 94.91 ± 2.86 | 93.75 ± 0.95 | 97.57 ± 0.66 |
| Electron balance (%) | 101.14 ± 0.70 | 95.35 ± 3.05 | 93.61 ± 0.44 | 97.41 ± 0.01 |

**Table S2.** Physiological parameters of S. cerevisiae JP1 during aerobic batch cultivations with either glucose, sucrose, an equimolar mixture of glucose and fructose, or fructose as sole carbon and energy source. All parameters were calculated for the exponential growth phase. The data represent the mean of two experiments and the average deviation.

| | **Strain: JP1** | | | | | | | | | | | |
|---|---|---|---|---|---|---|---|---|---|---|---|---|
| | Glucose | | | Sucrose | | | Glucose & Fructose | | | Fructose | | |
| $\mu_{MAX}$ ($h^{-1}$) | 0.28 | ± | 0.02 | 0.32 | ± | 0.00 | 0.30 | ± | 0.02 | 0.32 | ± | 0.02 |
| $Y_{X/S}$ ($g_{DM}$ $gGLC_{eq}^{-1}$) | 0.13 | ± | 0.01 | 0.14 | ± | 0.00 | 0.17 | ± | 0.01 | 0.15 | ± | 0.01 |
| $Y_{Ethanol/S}$ (g $gGLC_{eq}^{-1}$) | 0.42 | ± | 0.00 | 0.41 | ± | 0.00 | 0.44 | ± | 0.02 | 0.38 | ± | 0.00 |
| $Y_{CO2/S}$ (g $gGLC_{eq}^{-1}$) | 0.41 | ± | 0.01 | 0.43 | ± | 0.00 | 0.42 | ± | 0.03 | 0.42 | ± | 0.00 |
| $Y_{glycerol/S}$ (g $gGLC_{eq}^{-1}$) | 0.02 | ± | 0.00 | 0.02 | ± | 0.00 | 0.02 | ± | 0.00 | 0.02 | ± | 0.00 |
| $Y_{acetate/S}$ (g $gGLC_{eq}^{-1}$) | 0.00 | ± | 0.00 | 0.00 | ± | 0.00 | 0.00 | ± | 0.00 | 0.00 | ± | 0.00 |
| $Y_{succinate/S}$ (g $gGLC_{eq}^{-1}$) | 0.00 | ± | 0.00 | 0.00 | ± | 0.00 | 0.00 | ± | 0.00 | 0.00 | ± | 0.00 |
| $Y_{lactate/S}$ (g $gGLC_{eq}^{-1}$) | 0.00 | ± | 0.00 | 0.00 | ± | 0.00 | 0.00 | ± | 0.00 | 0.00 | ± | 0.00 |
| $qS_{MAX}$ ($mmolGLC_{eq}$ $g_{DM}^{-1}$ $h^{-1}$) | -11.67 | ± | 0.45 | -12.29 | ± | 0.12 | -10.08 | ± | 0.39 | -12.24 | ± | 0.07 |
| $qCO_2$ (mmol $g_{DM}^{-1}$ $h^{-1}$) | 19.75 | ± | 0.31 | 21.74 | ± | 0.19 | 17.53 | ± | 1.80 | 20.89 | ± | 0.05 |
| $qO_2$ (mmol $g_{DM}^{-1}$ $h^{-1}$) | -2.12 | ± | 0.09 | -2.19 | ± | 0.32 | -2.15 | ± | 0.31 | -2.43 | ± | 0.02 |
| RQ ($mmolCO_2$ $mmolO_2^{-1}$) | 9.31 | ± | 0.25 | 10.13 | ± | 1.40 | 8.19 | ± | 0.33 | 8.60 | ± | 0.04 |
| $q_{Ethanol}$ (mmol $g_{DM}^{-1}$ $h^{-1}$) | 19.23 | ± | 0.83 | 19.64 | ± | 0.26 | 17.18 | ± | 1.54 | 18.23 | ± | 0.14 |
| $q_{glycerol}$ (mmol $g_{DM}^{-1}$ $h^{-1}$) | 0.52 | ± | 0.08 | 0.48 | ± | 0.03 | 0.42 | ± | 0.00 | 0.39 | ± | 0.07 |
| $q_{acetate}$ (mmol $g_{DM}^{-1}$ $h^{-1}$) | 0.17 | ± | 0.09 | 0.08 | ± | 0.00 | 0.08 | ± | 0.00 | 0.10 | ± | 0.01 |
| $q_{succinate}$ (mmol $g_{DM}^{-1}$ $h^{-1}$) | 0.07 | ± | 0.00 | 0.06 | ± | 0.01 | 0.04 | ± | 0.02 | 0.05 | ± | 0.01 |
| $q_{lactate}$ (mmol $g_{DM}^{-1}$ $h^{-1}$) | 0.07 | ± | 0.01 | 0.05 | ± | 0.01 | 0.06 | ± | 0.00 | 0.07 | ± | 0.02 |
| Residual substrate ($gGLC_{eq}$ $l^{-1}$) | 0.73 | ± | 0.08 | 1.79 | ± | 0.15 | 0.90 | ± | 0.11 | 1.34 | ± | 0.10 |
| Carbon recovery (%) | 101.44 | ± | 1.35 | 101.59 | ± | 0.11 | 107.25 | ± | 5.47 | 97.12 | ± | 0.51 |
| Electron balance (%) | 104.06 | ± | 0.53 | 102.13 | ± | 0.49 | 110.76 | ± | 5.34 | 97.25 | ± | 0.66 |

**Table S3. .** Physiological parameters of *S. cerevisiae* UFMG-CM-Y259 during aerobic batch cultivations with either glucose, sucrose, an equimolar mixture of glucose and fructose, or fructose as sole carbon and energy source. All parameters were calculated for the exponential growth phase. The data represent the mean of two experiments and the average deviation.

| | **Strain: UFMG-CM-Y259** | | | | | | | | | | | |
|---|---|---|---|---|---|---|---|---|---|---|---|---|
| | Glucose | | | Sucrose | | | Glucose & Fructose | | | Fructose | | |
| $\mu_{MAX}$ ($h^{-1}$) | 0.29 | ± | 0.00 | 0.37 | ± | 0.01 | 0.35 | ± | 0.03 | 0.36 | ± | 0.02 |
| $Y_{X/S}$ ($g_{DM}$ $gGLC_{eq}^{-1}$) | 0.14 | ± | 0.00 | 0.16 | ± | 0.01 | 0.16 | ± | 0.01 | 0.16 | ± | 0.00 |
| $Y_{Ethanol/S}$ (g $gGLC_{eq}^{-1}$) | 0.40 | ± | 0.03 | 0.41 | ± | 0.01 | 0.38 | ± | 0.00 | 0.37 | ± | 0.00 |
| $Y_{CO2/S}$ (g $gGLC_{eq}^{-1}$) | 0.45 | ± | 0.02 | 0.43 | ± | 0.05 | 0.42 | ± | 0.01 | 0.41 | ± | 0.00 |
| $Y_{glycerol/S}$ (g $gGLC_{eq}^{-1}$) | 0.02 | ± | 0.00 | 0.02 | ± | 0.00 | 0.02 | ± | 0.00 | 0.03 | ± | 0.01 |
| $Y_{acetate/S}$ (g $gGLC_{eq}^{-1}$) | 0.01 | ± | 0.00 | 0.00 | ± | 0.00 | 0.01 | ± | 0.00 | 0.01 | ± | 0.00 |
| $Y_{succinate/S}$ (g $gGLC_{eq}^{-1}$) | 0.00 | ± | 0.00 | 0.00 | ± | 0.00 | 0.00 | ± | 0.00 | 0.00 | ± | 0.00 |
| $Y_{lactate/S}$ (g $gGLC_{eq}^{-1}$) | 0.00 | ± | 0.00 | 0.00 | ± | 0.00 | 0.00 | ± | 0.00 | 0.00 | ± | 0.00 |
| $qS_{MAX}$ ($mmolGLC_{eq}$ $g_{DM}^{-1}$ $h^{-1}$) | -11.96 | ± | 0.43 | -12.84 | ± | 0.21 | -12.19 | ± | 0.63 | -12.12 | ± | 0.84 |
| $qCO_2$ (mmol $g_{DM}^{-1}$ $h^{-1}$) | 20.77 | ± | 0.50 | 22.50 | ± | 2.00 | 20.96 | ± | 1.55 | 20.44 | ± | 1.39 |
| $qO_2$ (mmol $g_{DM}^{-1}$ $h^{-1}$) | -2.63 | ± | 0.31 | -3.34 | ± | 0.21 | -2.88 | ± | 0.18 | -3.03 | ± | 0.12 |
| RQ ($mmolCO_2$ $mmolO_2^{-1}$) | 8.01 | ± | 1.12 | 6.81 | ± | 1.03 | 7.34 | ± | 1.00 | 6.73 | ± | 0.20 |
| $q_{Ethanol}$ (mmol $g_{DM}^{-1}$ $h^{-1}$) | 17.63 | ± | 0.76 | 20.82 | ± | 0.65 | 18.00 | ± | 1.06 | 17.57 | ± | 1.26 |
| $q_{glycerol}$ (mmol $g_{DM}^{-1}$ $h^{-1}$) | 0.47 | ± | 0.07 | 0.48 | ± | 0.05 | 0.40 | ± | 0.07 | 0.60 | ± | 0.13 |
| $q_{acetate}$ (mmol $g_{DM}^{-1}$ $h^{-1}$) | 0.21 | ± | 0.03 | 0.15 | ± | 0.01 | 0.20 | ± | 0.03 | 0.30 | ± | 0.07 |
| $q_{succinate}$ (mmol $g_{DM}^{-1}$ $h^{-1}$) | 0.05 | ± | 0.00 | 0.05 | ± | 0.00 | 0.08 | ± | 0.03 | 0.07 | ± | 0.00 |
| $q_{lactate}$ (mmol $g_{DM}^{-1}$ $h^{-1}$) | 0.10 | ± | 0.02 | 0.12 | ± | 0.01 | 0.12 | ± | 0.03 | 0.09 | ± | 0.00 |
| Residual substrate ($gGLC_{eq}$ $l^{-1}$) | 0.03 | ± | 0.03 | 1.03 | ± | 0.84 | 0.91 | ± | 0.40 | 2.28 | ± | 0.19 |
| Carbon recovery (%) | 103.25 | ± | 6.11 | 104.55 | ± | 2.95 | 98.80 | ± | 1.74 | 99.04 | ± | 0.94 |
| Electron balance (%) | 99.05 | ± | 2.05 | 107.25 | ± | 0.76 | 99.09 | ± | 0.96 | 100.00 | ± | 0.96 |

**Table S4.** Point mutations in the *SUC2* sequence of *S. cerevisiae* strains JP1 and UFMG-CM-Y259 with respect to the reference strain S288C. Nucleotide and aminoacid (aa) changes are indicated in red and blue, respectively. Colored in grey are mutations shared by the two strains.

| **Gene:** *SUC2* **Accession number:** V01311 | | | | | |
|---|---|---|---|---|---|
| | | **Mutation** | | | |
| **Position (bp)** | **Original codon** | **aa** | **Mutated codon** | **aa** | **Type** |
| **JP1** | | | | | |
| 228 | GCT | A | ACT | T | Missense |
| 249 | AAT | N | CAT | H | Missense |
| 261 | CAA | Q | GAA | E | Missense |
| 338 | ACG | T | ACC | T | Silent |
| 386 | CGC | A | GCA | A | Silent |
| 437 | TCT | S | TCC | S | Silent |
| 599 | TCT | S | TCG | S | Silent |
| 621 | CTA | L | TTA | L | Silent |
| 641 | AAC | N | AAT | N | Silent |
| 665 | TAC | Y | TAT | Y | Silent |
| 716 | TCT | S | TCC | S | Silent |
| 866 | TTG | L | TTA | L | Silent |
| 1082 | GGT | G | GGC | G | Silent |
| 1085 | CCC | P | CCA | P | Silent |
| 1103 | ACT | T | ACC | T | Silent |
| 1190 | GTT | V | GTC | V | Silent |
| 1223 | TTT | F | TTC | F | Silent |
| 1224 | GCC | A | CCC | P | Missense |
| 1544 | ACT | T | ACC | T | Silent |
| **UFMG-CM-Y259** | | | | | |
| 135 | TTG | L | CTG | L | Silent |
| 249 | AAT | N | CAT | H | Missense |
| 261 | CAA | Q | GAA | E | Missense |
| 338 | ACG | T | ACC | T | Silent |
| 386 | GCG | A | GCA | A | Silent |
| 437 | TCT | S | TCC | S | Silent |
| 473 | AAG | K | AAA | K | Silent |
| 641 | AAC | N | AAT | N | Silent |
| 1082 | GGT | G | GGC | G | Silent |
| 1103 | ACT | T | ACC | T | Silent |
| 1334 | GTC | V | GTT | V | Silent |

| Strain | | *SUC2* sequence | |
|---|---|---|---|
| JP1 | 1 | MLLQAFLFLLAGFAAKISASMTNETSDRPLVHFTPNKGWMNDPNGLWYDE | 50 |
| UFMG-CM-Y259 | | MLLQAFLFLLAGFAAKISASMTNETSDRPLVHFTPNKGWMNDPNGLWYDE | |
| CEN.PK113-7D | | MLLQAFLFLLAGFAAKISASMTNETSDRPLVHFTPNKGWMNDPNGLWYDE | |
| S288C | | MLLQAFLFLLAGFAAKISASMTNETSDRPLVHFTPNKGWMNDPNGLWYDE | |
| | 51 | KDAKWHLYFQYNPNDTVWGTPLFWGHTTSDDLTHWEDEPIAIAPKRNDSG | 100 |
| | | KDAKWHLYFQYNPNDTVWGTPLFWGHATSDDLTHWEDEPIAIAPKRNDSG | |
| | | KDAKWHLYFQYNPNDTVWGTPLFWGHATSDDLTNWEDQPIAIAPKRNDSG | |
| | | KDAKWHLYFQYNPNDTVWGTPLFWGHATSDDLTNWEDQPIAIAPKRNDSG | |
| | 101 | AFSGSMVVDYNNTSGFFNDTIDPRQRCVAIWTYNTPESEEQYISYSLDGG | 150 |
| | | AFSGSMVVDYNNTSGFFNDTIDPRQRCVAIWTYNTPESEEQYISYSLDGG | |
| | | AFSGSMVVDYNNTSGFFNDTIDPRQRCVAIWTYNTPESEEQYISYSLDGG | |
| | | AFSGSMVVDYNNTSGFFNDTIDPRQRCVAIWTYNTPESEEQYISYSLDGG | |
| | 151 | YTFTEYQKNPVLAANSTQFRDPKVFWYEPSQKWIMTAAKSQDYKIEIYSS | 200 |
| | | YTFTEYQKNPVLAANSTQFRDPKVFWYEPSQKWIMTAAKSQDYKIEIYSS | |
| | | YTFTEYQKNPVLAANSTQFRDPKVFWYEPSQKWIMTAAKSQDYKIEIYSS | |
| | | YTFTEYQKNPVLAANSTQFRDPKVFWYEPSQKWIMTAAKSQDYKIEIYSS | |
| | 201 | DDLKSWKLESAFANEGFLGYQYECPGLIEVPTEQDPSKSYWVMFISINPG | 250 |
| | | DDLKSWKLESAFANEGFLGYQYECPGLIEVPTEQDPSKSYWVMFISINPG | |
| | | DDLKSWKLESAFANEGFLGYQYECPGLIEVPTEQDPSKSYWVMFISINPG | |
| | | DDLKSWKLESAFANEGFLGYQYECPGLIEVPTEQDPSKSYWVMFISINPG | |
| | 251 | APAGGSFNQYFVGSFNGTHFEAFDNQSRVVDFGKDYYALQTFFNTDPTYG | 300 |
| | | APAGGSFNQYFVGSFNGTHFEAFDNQSRVVDFGKDYYALQTFFNTDPTYG | |
| | | APAGGSFNQYFVGSFNGTHFEAFDNQSRVVDFGKDYYALQTFFNTDPTYG | |
| | | APAGGSFNQYFVGSFNGTHFEAFDNQSRVVDFGKDYYALQTFFNTDPTYG | |
| | 301 | SALGIAWASNWEYSAFVPTNPWRSSMSLVRKFSLNTEYQANPETELINLK | 350 |
| | | SALGIAWASNWEYSAFVPTNPWRSSMSLVRKFSLNTEYQANPETELINLK | |
| | | SALGIAWASNWEYSAFVPTNPWRSSMSLVRKFSLNTEYQANPETELINLK | |
| | | SALGIAWASNWEYSAFVPTNPWRSSMSLVRKFSLNTEYQANPETELINLK | |
| | 351 | AEPILNISNAGPWSRFATNTTLTKANSYNVDLSNSTGTLEFELVYAVNTT | 400 |
| | | AEPILNISNAGPWSRFATNTTLTKANSYNVDLSNSTGTLEFELVYAVNTT | |
| | | AEPILNISNAGPWSRFATNTTLTKANSYNVDLSNSTGTLEFELVYAVNTT | |
| | | AEPILNISNAGPWSRFATNTTLTKANSYNVDLSNSTGTLEFELVYAVNTT | |
| | 401 | QTISKSVFPDLSLWFKGLEDPEEYLRMGFEVSASSFFLDRGNSKVKFVKE | 450 |
| | | QTISKSVFADLSLWFKGLEDPEEYLRMGFEVSASSFFLDRGNSKVKFVKE | |
| | | QTISKSVFADLSLWFKGLEDPEEYLRMGFEVSASSFFLDRGNSKVKFVKE | |
| | | QTISKSVFADLSLWFKGLEDPEEYLRMGFEVSASSFFLDRGNSKVKFVKE | |
| | 451 | NPYFTNRMSVNNQPFKSENDLSYYKVYGLLDQNILELYFNDGDVVSTNTY | 500 |
| | | NPYFTNRMSVNNQPFKSENDLSYYKVYGLLDQNILELYFNDGDVVSTNTY | |
| | | NPYFTNRMSVNNQPFKSENDLSYYKVYGLLDQNILELYFNDGDVVSTNTY | |
| | | NPYFTNRMSVNNQPFKSENDLSYYKVYGLLDQNILELYFNDGDVVSTNTY | |
| | 501 | FMTTGNALGSVNMTTGVDNLFYIDKFQVREVK | 532 |
| | | FMTTGNALGSVNMTTGVDNLFYIDKFQVREVK | |
| | | FMTTGNALGSVNMTTGVDNLFYIDKFQVREVK | |
| | | FMTTGNALGSVNMTTGVDNLFYIDKFQVREVK | |

**Figure S6.** Alignment of the translated *SUC2* sequence of *S. cerevisiae* CEN.PK113-7D, JP1, UFMG-CM-Y259, and S288C (reference). Missense mutations in at least one strain are highlighted in yellow. The alignment was performed using the Basic Local Alignment Search Tool with a translated nucleotide query (BLASTX) (Altschul et al. 1997).

**Table S5.** Point mutations within 1000 bp upstream of the *SUC2* ORF of strains JP1 and UFMG-CM-Y259 with respect to the reference strain S288C. Colored in grey are mutations shared by the two strains.

| Position (bp) | nucleotide mutation | | |
|---|---|---|---|
| **JP1** | | | |
| -929 | C | → | A |
| -805 | A | → | C |
| -776 | C | → | T |
| -704 | C | → | T |
| -698 | A | → | G |
| -184 | C | → | CTTTT, CTTTTTTTTT |
| -86 | CT | → | C |
| **UFMG-CM-Y259** | | | |
| -971 | GT | → | G |
| -805 | A | → | C |
| -776 | C | → | T |
| -737 | G | → | A |
| -317 | T | → | C |
| -236 | A | → | T |
| -184 | C | → | CTTTT, CTTTTTTTTT |
| -170 | A | → | T |

**Table S6.** Point mutations in the *AGT1* sequence of *S. cerevisiae* strains JP1 and UFMG-CM-Y259 with respect to the reference strain CAT-1 (S288C does not have a functional *AGT1* allele). Nucleotide and aminoacid (aa) changes are indicated in red and blue, respectively. Colored in grey are mutations shared by the two strains.

| **Gene:** *AGT1* **Accession number:** MF374788 | | | | | |
|---|---|---|---|---|---|
| | | **Mutation** | | | |
| **Position (bp)** | **Original codon** | **aa** | **Mutated codon** | **aa** | **Type** |
| | | **JP1** | | | |
| 118 | ATT | N | GAT | D | Missense |
| 233 | ACG | T | ATG | M | Missense |
| 305 | ATA | I | AAA | K | Missense |
| 383 | AAC | N | AGC | S | Missense |
| 489 | GTC | V | GTT | V | Silent |
| 491<br>492 | CAA | Q | CTT | L | Missense |
| 523 | CCT | P | ACT | T | Missense |
| 634 | GTG | V | ATG (80%) | M | Missense |
| 675 | CAG | Q | CAA | Q | Silent |
| 676 | GGT | G | AGT (68%)<br>TGT (32%) | S<br>C | Missense<br>Missense |
| 682<br>684 | ACT | T | GCT (69%)*<br>ACC (27%)* | A<br>T | Missense<br>Silent |
| 688 | ACT | T | TCT (25%) | S | Missense |
| 819 | TCT | S | TCC | S | Silent |
| 997 | ATT | I | GTT | V | Missense |
| 1024 | TTG | L | ATG | M | Missense |
| 1075 | GAT | D | AAT | N | Missense |
| 1123 | GCT | A | ACT | T | Missense |
| 1142 | ACT | T | AGT | S | Missense |
| 1153<br>1154 | TGT | C | GTT | V | Missense |
| 1225 | GTA | V | CTA | L | Missense |
| 1336 | TTA | L | CTA | L | Silent |
| 1342 | GTT | V | ATT | I | Missense |
| 1375 | GGC | G | AGC | S | Missense |
| 1461 | GTA | V | GTT | V | Silent |
| 1462 | ACT | T | GCT | A | Missense |
| 1525 | ATC | I | CTC | L | Missense |
| 1539 | ATC | I | ATT | I | Silent |
| 1667 | AGT | S | ACT | T | Missense |
| | | **UFMG-CM-Y259** | | | |

| Gene: *AGT1* Accession number: MF374788 | | | | | |
|---|---|---|---|---|---|
| | | | Mutation | | |
| Position (bp) | Original codon | aa | Mutated codon | aa | Type |
| 118 | ATT | N | GAT | D | Missense |
| 233 | ACG | T | ATG | M | Missense |
| 288 | TTA | L | TTG | L | Silent |
| 305 | ATA | I | AAA | K | Missense |
| 383 | AAC | N | AGC | S | Missense |
| 491<br>492 | CAA | Q | CTT | L | Missense |
| 523 | CCT | P | ACT | T | Missense |
| 675 | CAG | Q | CAA | Q | Silent |
| 676 | GGT | G | AGT (70%)<br>TGT (30%) | S<br>C | Missense<br>Missense |
| 682<br>684 | ACT | T | GCT (71%)<br>ACC (29%) | A<br>T | Missense<br>Silent |
| 688 | ACT | T | TCT (26%) | S | Missense |
| 819 | TCT | S | TCC | S | Silent |
| 997 | ATT | I | GTT | V | Missense |
| 1075 | GAT | D | AAT | N | Missense |
| 1123 | GCT | A | ACT | T | Missense |
| 1142 | ACT | T | AGT | S | Missense |
| 1153 | TGT | C | GTT | V | Missense |
| 1225 | GTA | V | CTA | L | Missense |
| 1342<br>1344 | GTT | V | ATC | I | Missense |
| 1375 | GGC | G | AGC | S | Missense |
| 1462 | ACT | T | GCT | A | Missense |
| 1525 | ATC | I | CTC | L | Missense |
| 1667 | AGT | S | ACT | T | Missense |

| Strain | | *AGT1* sequence | |
|---|---|---|---|
| JP1 | 1 | MKNIISLVSKKKAASKNEDKNISESSRDIVNQQEVFNTEDFEEGKKDSAFELDHLEFTTNSAQLGDSDEDNENVINEMNATDDANEANSEEKSMTLKQAL | 100 |
| UFMG-CM-Y259 | | MKNIISLVSKKKAASKNEDKNISESSRDIVNQQEVFNTEDFEEGKKDSAFELDHLEFTTNSAQLGDSDEDNENVINEMNATDDANEANSEEKSMTLKQAL | |
| CAT-1 | | MKNIISLVSKKKAASKNEDKNISESSRDIVNQQEVFNTENFEEGKKDSAFELDHLEFTTNSAQLGDSDEDNENVINETNATDDANEANSEEKSMTLKQAL | |
| | 101 | LKYPKAALWSILVSTTLVMEGYDTALLSALYALPVFQRKFGTLNGEGSYEITSQWQIGLNMCVLCGEMIGLQITTYMVEFMGNRYTMITALGLLTAYIFI | 200 |
| | | LKYPKAALWSILVSTTLVMEGYDTALLSALYALPVFQRKFGTLNGEGSYEITSQWQIGLNMCVLCGEMIGLQITTYMVEFMGNRYTMITALGLLTAYIFI | |
| | | LIYPKAALWSILVSTTLVMEGYDTALLNALYALPVFQRKFGTLNGEGSYEITSQWQIGLNMCVQCGEMIGLQITPYMVEFMGNRYTMITALGLLTAYIFI | |
| | 201 | LYYCKSLAMIAMGQVLSAMPWGCFQS/CLAVSYASEVCPLALRYYMTSYSNICWLFGQIFASGIMKNSQENLGNSDLGYKLPFALQWIWPAPLMIGIFFAPE | 300 |
| | | LYYCKSLAMIAVGQVLSAMPWGCFQS/CLAVSYASEVCPLALRYYMTSYSNICWLFGQIFASGIMKNSQENLGNSDLGYKLPFALQWIWPAPLMIGIFFAPE | |
| | | LYYCKSLAMIAVGQVLSAMPWGCFQGLTVTYASEVCPLALRYYMTSYSNICWLFGQIFASGIMKNSQENLGNSDLGYKLPFALQWIWPAPLMIGIFFAPE | |
| | 301 | SPWWLVRKDRVAEARKSLSRILSGKGAEKDIQVDLTLKQIEMTIEKERLLASKSGSFFNCFKGVNGRRTRLACLTWVAQNSSGAVLLGYSTYFFERAGMA | 400 |
| | | SPWWLVRKDRVAEARKSLSRILSGKGAEKDIQVDLTLKQIELTIEKERLLASKSGSFFNCFKGVNGRRTRLACLTWVAQNSSGAVLLGYSTYFFERAGMA | |
| | | SPWWLVRKDRVAEARKSLSRILSGKGAEKDIQIDLTLKQIELTIEKERLLASKSGSFFDCFKGVNGRRTRLACLAWVAQNTSGACLLGYSTYFFERAGMA | |
| | 401 | TDKAFTFSLIQYCLGLAGTLCSWVISGRVGRWTILTYGLAFQMVCLFIIGGMGFGSGSSASNGAGGIIIAISFFYNAGIGAVVYCIVAEIPSAELRTKTI | 500 |
| | | TDKAFTFSLIQYCLGLAGTLCSWVISGRVGRWTILTYGLAFQMVCLFIIGGMGFGSGSSASNGAGGIIIAISFFYNAGIGAVVYCIVAEIPSAELRTKTI | |
| | | TDKAFTFSVIQYCLGLAGTLCSWVISGRVGRWTILTYGLAFQMVCLFVIGGMGFGSGSGASNGAGGIIIAISFFYNAGIGAVVYCIVTEIPSAELRTKTI | |
| | 501 | VLARICYNLMAVINAILTPYMLNVSDWNWGAKTGLYWGGFTAVTLAWVIIDLPETTGRTFSEINELFNQGVPARKFASTVVDPFGKGKTQHDSLADESIS | 600 |
| | | VLARICYNLMAVINAILTPYMLNVSDWNWGAKTGLYWGGFTAVTLAWVIIDLPETTGRTFSEINELFNQGVPARKFASTVVDPFGKGKTQHDSLADESIS | |
| | | VLARICYNIMAVINAILTPYMLNVSDWNWGAKTGLYWGGFTAVTLAWVIIDLPETSGRTFSEINELFNQGVPARKFASTVVDPFGKGKTQHDSLADESIS | |
| | 601 | QSSSIKQRELNAADKC | 616 |
| | | QSSSIKQRELNAADKC | |
| | | QSSSIKQRELNAADKC | |

**Figure S7.** Alignment of the translated *AGT1* sequence of *S. cerevisiae* JP1 and UFMG-CM-Y259, and CAT-1 (reference). Missense mutations in at least one strain are highlighted in yellow. The alignment was performed using the Basic Local Alignment Search Tool with a translated nucleotide query (BLASTX) (Altschul et al. 1997).

**Table S7.** Point mutations in the *HXT3* sequence of *S. cerevisiae* strains JP1 and UFMG-CM-Y259 with respect to the reference strain S288C. Nucleotide and aminoacid (aa) changes are indicated in red and blue, respectively. Colored in grey are mutations shared by the two strains.

| Gene: *HXT3* Accession number: | | | | | |
|---|---|---|---|---|---|
| | | Mutation | | | |
| **Position (bp)** | **Original codon** | **aa** | **Mutate codon** | **aa** | **Type** |
| **JP1** | | | | | |
| 501 | TAT | Y | TAC | Y | Silent |
| 906 | GCT | A | GCC | A | Silent |
| 913<br>915 | TCA | S | ACT | T | Missense |
| 921 | TCA | S | TCT | S | Silent |
| 927 | GGT | G | GGC | G | Silent |
| 930 | GAG | E | GAA | E | Silent |
| 933 | TTG | L | TTA | L | Silent |
| 945 | AAG | K | AAA | K | Silent |
| 948 | CCG | P | CCA | P | Silent |
| 958<br>960 | AAG | K | CAA | Q | Missense |
| 1443 | CCA | P | CCG | P | Silent |
| **UFMG-CM-Y259** | | | | | |
| 36 | AAG | K | AAA | K | Silent |
| 148 | ACC | T | GCC | A | Missense |
| 159 | AAT | N | AAC | N | Silent |
| 174 | GCA | A | GCC | A | Silent |
| 393 | GCT | A | GCC | A | Silent |
| 414 | GGT | G | GGA | G | Silent |
| 480 | TCC | S | TCT | S | Silent |
| 501 | TAT | Y | TAC | Y | Silent |
| 867 | GAG | E | GAA | E | Silent |
| 948 | CCG | P | CCA | P | Silent |
| 1159 | TTG | L | CTG | L | Silent |
| 1174 | ATT | I | GTT | V | Missense |
| 1225 | CTA | L | TTA | L | Silent |
| 1282 | GTC → TC | | | | Deletion |
| 1284 | GTC | V | TGT | C | Missense |
| 1288 | GC → CGC | | | | Insertion |
| 1291 | GCC | A | GCA | A | Silent |
| 1452 | ACT | T | ACC | T | Silent |

| Strain | | *HXT3* sequence | |
|---|---|---|---|
| JP1 | 1 | MNSTPDLISPQKSSENSNADLPSNSSQVMNMPEEKGVQDDFQAEADQVLTNPNTGKGAYVTVSICCVMVAFGGFVFGWDTGTISGFVAQTDFLRRFGMKH | 100 |
| UFMG-CM-Y259 | | MNSTPDLISPQKSSENSNADLPSNSSQVMNMPEEKGVQDDFQAEADQVLANPNTGKGAYVTVSICCVMVAFGGFVFGWDTGTISGFVAQTDFLRRFGMKH | |
| CEN.PK113-7D | | MNSTPDLISPQKSSENSNADLPSNSSQVMNMPEEKGVQDDFQAEADQVLTNPNTGKGAYVTVSICCVMVAFGGFVFGWDTGTISGFVAQTDFLRRFGMKH | |
| S288C | | MNSTPDLISPQKSSENSNADLPSNSSQVMNMPEEKGVQDDFQAEADQVLTNPNTGKGAYVTVSICCVMVAFGGFVFGWDTGTISGFVAQTDFLRRFGMKH | |
| | 101 | KDGSYYLSKVRTGLIVSIFNIGCAIGGIILAKLGDMYGRKMGLIVVVVIYIIGIIIQIASINKWYQYFIGRIISGLGVGGIAVLSPMLISEVAPKEMRGT | 200 |
| | | KDGSYYLSKVRTGLIVSIFNIGCAIGGIILAKLGDMYGRKMGLIVVVVIYIIGIIIQIASINKWYQYFIGRIISGLGVGGIAVLSPMLISEVAPKEMRGT | |
| | | KDGSYYLSKVRTGLIVSIFNIGCAIGGIILAKLGDMYGRKMGLIVVVVIYIIGIIIQIASINKWYQYFIGRIISGLGVGGIAVLSPMLISEVAPKEMRGT | |
| | | KDGSYYLSKVRTGLIVSIFNIGCAIGGIILAKLGDMYGRKMGLIVVVVIYIIGIIIQIASINKWYQYFIGRIISGLGVGGIAVLSPMLISEVAPKEMRGT | |
| | 201 | LVSCYQLMITLGIFLGYCTNFGTKNYSNSVQWRVPLGLCFAWALFMIGGMTFVPESPRYLVEAGQIDEARASLSKVNKVAPDHPFIQQELEVIEASVEEA | 300 |
| | | LVSCYQLMITLGIFLGYCTNFGTKNYSNSVQWRVPLGLCFAWALFMIGGMTFVPESPRYLVEAGQIDEARASLSKVNKVAPDHPFIQQELEVIEASVEEA | |
| | | LVSCYQLMITLGIFLGYCTNFGTKNYSNSVQWRVPLGLCFAWALFMIGGMTFVPESPRYLVEAGQIDEARASLSKVNKVAPDHPFIQQELEVIEASVEEA | |
| | | LVSCYQLMITLGIFLGYCTNFGTKNYSNSVQWRVPLGLCFAWALFMIGGMTFVPESPRYLVEAGQIDEARASLSKVNKVAPDHPFIQQELEVIEASVEEA | |
| | 301 | RAAGTASWGELFTGKPAMFQRTMMGIMIQSLQQLTGDNYFFYYGTTVFNAVGMSDSFETSIVFGVVNFFSTCCSLYTVDRFGRRNCLLYGAIGMVCCYVV | 400 |
| | | RAAGSASWGELFTGKPAMFKRTMMGIMIQSLQQLTGDNYFFYYGTTVFNAVGMSDSFETSIVFGVVNFFSTCCSLYTVDRFGRRNCLLYGAVGMVCCYVV | |
| | | RAAGSASWGELFTGKPAMFKRTMMGIMIQSLQQLTGDNYFFYYGTTVFNAVGMSDSFETSIVFGVVNFFSTCCSLYTVDRFGRRNCLLYGAIGMVCCYVV | |
| | | RAAGSASWGELFTGKPAMFKRTMMGIMIQSLQQLTGDNYFFYYGTTVFNAVGMSDSFETSIVFGVVNFFSTCCSLYTVDRFGRRNCLLYGAIGMVCCYVV | |
| | 401 | YASVGVTRLWPNGEGNGSSKGAGNCMIVFACFYIFCFATTWAPIAYVVISETFPLRVKSKAMSIATAANWLWGFLIGFFTPFITGAINFYYGYVFMGCMV | 500 |
| | | YASVGVTRLWPNGEGNGSSKGAGNCMICFACFYIFCFATTWAPIAYVVISETFPLRVKSKAMSIATAANWLWGFLIGFFTPFITGAINFYYGYVFMGCMV | |
| | | YASVGVTRLWPNGEGNGSSKGAGNCMIVFACFYIFCFATTWAPIAYVVISETFPLRVKSKAMSIATAANWLWGFLIGFFTPFITGAINFYYGYVFMGCMV | |
| | | YASVGVTRLWPNGEGNGSSKGAGNCMIVFACFYIFCFATTWAPIAYVVISETFPLRVKSKAMSIATAANWLWGFLIGFFTPFITGAINFYYGYVFMGCMV | |
| | 501 | FAYFYVFFFVPETKGLTLEEVNDMYAEGVLPWKSASWVPTSQRGANYDADALMHDDQPFYKKMFGKK | 567 |
| | | FAYFYVFFFVPETKGLTLEEVNDMYAEGVLPWKSASWVPTSQRGANYDADALMHDDQPFYKKMFGKK | |
| | | FAYFYVFFFVPETKGLTLEEVNDMYAEGVLPWKSASWVPTSQRGANYDADALMHDDQPFYKKMFGKK | |
| | | FAYFYVFFFVPETKGLTLEEVNDMYAEGVLPWKSASWVPTSQRGANYDADALMHDDQPFYKKMFGKK | |

**Figure S8.** Alignment of the translated *HXT3* sequence of *S. cerevisiae* CEN.PK113-7D, JP1, UFMG-CM-Y259, and S288C (reference). Missense mutations in at least one strain are highlighted in yellow. The alignment was performed using the Basic Local Alignment Search Tool with a translated nucleotide query (BLASTX) (Altschul et al. 1997).

**Table S9.** Point mutations within 1000 bp upstream of the *HXT3* ORF of *S. cerevisiae* strains JP1 and UFMG-CM-Y259 with respect to the reference strain S288C. Only one mutation is shared by the two strains (in gray color).

| Position (bp) | nucleotide mutation | | |
|---|---|---|---|
| **JP1** | | | |
| -118 | G | → | A |
| -601 | A | → | T |
| **UFMG-CM-Y259** | | | |
| -65 | T | → | C |
| -124 | G | → | A |
| -125 | CT | → | C |
| -140 | GA | → | G |

| Position (bp) | nucleotide mutation | | |
| --- | --- | --- | --- |
| -519 | T | → | C |
| -586 | C | → | G |
| -601 | A | → | T |
| -689 | T | → | TAA |
| -820 | A | → | AGC |
| -858 | C | → | T |
| -904 | C | → | T |
| -925 | T | → | C |
| -926 | C | → | G |
